# Caspase Cleavage of Kaposi Sarcoma-Associated Herpesvirus Proteins: A role for K5 in Preventing Caspase-Mediated Cell Death during Lytic Replication

**DOI:** 10.1101/2025.04.08.647834

**Authors:** Yana Astter, David A. Davis, Emma Treco, Prabha Shrestha, Alexandra Stream, Muzammel Haque, Naomi Mulugeta, Robert Yarchoan

## Abstract

We previously reported that Kaposi sarcoma-associated herpesvirus (KSHV) latency-associated nuclear antigen (LANA) acts as a pseudo-substrate for caspases-1 and 3, thereby interfering with their inflammatory and apoptotic activity, respectively. To determine if other KSHV proteins undergo caspase cleavage, we screened the KSHV proteome for potential caspase cleavage sites. Using SitePrediction (SP), 30 KSHV proteins with potential caspase-cleavage sites were identified. Among those with highest SP score was an early lytic protein, K5. Treatment of BJAB K5-FLAG cells with ⍺Fas, an apoptotic stimulus, led to caspase-processing of full length K5-FLAG and generation of a C-terminal peptide fragment. Using mass spectrometry, we determined that K5-FLAG undergoes caspase cleavage at D222. K5 was also cleaved by caspases in KSHV infected cells induced to lytic replication. Importantly, the expression of K5-FLAG significantly inhibited ⍺Fas-induced caspase-mediated cell death. To determine if K5 plays a protective role in KSHV infected cells, iSLKK cells infected with wild type or K5 knockout BAC16 virus were induce to lytic replication to activate caspases. Although lytic induction showed little effect on the viability of WT infected cells, the viability of K5-knockout cells decreased by 25%. Thus, K5 may protect KSHV-infected cells from caspase-mediated cell death during lytic replication. Interestingly, cleavage of K5 by caspases did not affect its ability to downregulate MHC-1 surface expression. Overall, these data suggest that K5 not only downregulates immunologic surface marker expression to avoid immune recognition but may also play an additional role in mitigating caspase-mediated cell death during KSHV lytic replication.

## Introduction

Most viruses must overcome one of the most fundamental innate cellular defense responses: the induction of caspase-mediated cell death that destroys infected cells before the virus can replicate. Viruses that infect eukaryotic cells have evolved means to prevent, inhibit, or redirect caspase activation upon viral infection. An early example of this was p35 protein from baculovirus which irreversibly inhibits caspases (1, 2) (for review see (3)). For chronic viruses like Kaposi sarcoma-associated herpesvirus (KSHV) or Epstein Barr virus (EBV), mechanisms to thwart caspase activation are essential to allow the establishment of latency (3–5). KSHV, which is the cause of Kaposi sarcoma (KS), primary effusion lymphoma (PEL), multicentric Castleman’s disease (MCD) and KSHV inflammatory cytokine syndrome (KICS), has evolved mechanisms to deal with multiple aspects of caspase activation. For example, vFLIP protein of KSHV can prevent caspase-8 activation by preventing the death receptor complex from activating caspase-8 (6–8).

We previously showed that KSHV latency-associated nuclear antigen (LANA) encodes two caspase cleavage sites, one in the N-terminus and one in the C-terminus (9). These sites may act as decoy substrates for caspase-3 and caspase-1, aiding in the prevention of apoptosis and activation of the inflammasome, respectively. The ORF63 lytic protein from KSHV acts as a viral homolog to NLRP1 and blocks NLRP1-mediated immune responses including caspase-1 activation and IL-1β and IL-18 production (10).

While inhibition of caspases by KSHV generally serves to allow lytic replication by preventing cell death, there are complexities to these interactions. For example, the increase in KSHV-induced caspase activity may benefit KSHV infection by antagonizing type I interferon responses (11, 12). Also, caspase-7 cleavage of the early lytic protein ORF57 (mRNA transcript accumulation [MTA]) by caspase-7 appears to attenuate the level of virus production; this allows for optimal conditions during viral release that could aid in immune avoidance and increase virus dissemination (13). However, in general, it’s more important for viral infection to limit caspase activity.

To follow up on previous studies of the caspase cleavage of LANA and ORF57 by caspases, we investigated the degree to which other KSHV-encoded proteins might be cleaved by caspases and the effects resulting from these interactions. We used SitePrediction (SP) to predict potential caspase cleavage sites in KSHV-encoded proteins. To our surprise, SP predicted that many KSHV proteins contain likely caspase cleavage sites. One particularly high scoring site was identified in the early lytic protein, K5. We explored this further to confirm whether K5 is cleaved by caspases in KSHV-infected cells and if so, to determine what role this cleavage might play in KSHV pathogenesis. K5 was cleaved by caspases-3 and 8 in infected cells and mechanistic studies suggest it can blunt cell death mediated by caspase-8 activation.

## Results

### Identification of predicted caspase-cleavage sites in KSHV-encoded proteins using SitePrediction

We used SitePrediction (https://www.irc.ugent.be/prx/bioit2-public/SitePrediction/reference.php) (14) to screen 87 KSHV-encoded proteins for potential caspase cleavage sites and the class of caspases predicted to cleave these sites. Sixty-six of the KSHV proteins had SP scores, ranging from a low of 25 to a high of 2787 (Table S1), predicting that they might be cleaved by caspases to varying degrees (the higher the score the more likely it might be cleaved by caspases). An additional 21 KSHV proteins (K1, K4, K4.1, K4.2, K6, K7, K9, K12a, ORF11, ORF16/vBcl2, ORF18, ORF29a, ORF36, ORF41, ORF47, ORF52, ORF53, ORF58, ORF65, ORF66, and ORF69) were negative for predicted caspase cleavage sites (Table S1). Using a SP cutoff score of 100, there were 35 KSHV proteins with predicted caspase cleavage sites, and 13 of these had a score of 500 or higher, indicating a high likelihood of cleavage (Table 1). Table 1 lists from highest to lowest the top scoring predicted cleavage sites found within the KSHV proteins that received a score over 100. Eighteen of the 35 proteins contained multiple predicted sites with a score over 100, all of which are indicated in Table 1 (Table 1). Among these 35 proteins with sites scoring over 100 were ORF57/MTA and ORF73/LANA, two viral proteins which have already been reported to be cleaved by caspases *in vitro* and in KSHV-infected cells(9, 13), providing some confidence in the use of SP to identify additional caspase cleavage sites in KSHV proteins. This said, while we had previously confirmed two caspase cleavage sites in LANA, one close to the N-terminus DSVD^53^*GREC and the other near the C-terminus,MEVD^906^*YPVV (9), only the N-terminus cleavage site had a score >100 (score 687); the C-terminal site had a SP score of 28 for caspase-3. Thus, while these programs can be a useful guide for prioritizing additional KSHV proteins with caspase cleavage sites, some sites could be missed, and not all predicted sites will end up being bona fide sites. The caspase cleavage site in ORF57 had the 4^th^ highest SP score (1742) while the N-terminal site in LANA had the 11^th^ highest SP score (687). The top scoring site predicted to be cleaved by caspases was within the ORF45 tegument protein with a score of 2787 for caspase processing predicted at D99 (Table 1). Sites within two other KSHV proteins K10.5/vIRF3 and K5/MIR2 had the second and third highest SP scores, respectively. Interestingly, 32 of the 35 proteins with scores over 100 are predicted by SP to be cleaved by executioner caspases (caspases-3,6,7) although sites for initiator (caspase-8) and inflammatory (caspase-1) are also represented (Table 1). Also, certain sites are predicted to be cleaved by multiple caspases. For example, the N-terminal cleavage site in LANA is predicted to be cleaved by initiator and executioner caspases. The high scoring site in K5 is predicted to be cleaved by all classes of caspases (Table 1).

**Table 1:** identification of KSHV Proteins with Potential Caspase-Cleavage sites using SitePrediction. KSHV protein sequences translated from the KSHV DNA sequence from BCBL-1 (accession number U93872.2) were analyzed for caspase cleavage sites for caspases-1,3,6,7, and 8 using site prediction (SP) (https://www.irc.ugent.be/prx/bioit2-public/SitePrediction/reference.php). The top scoring site scoring over 100 for each protein is shown in column 2.Column 3 shows the score and the class of caspases ((initiator caspase-8 (In), inflammatory caspase-1 (Inf), executioner caspases-3,6,7 (Ex)) predicted to cut that site. An * indicates that this protein has one or more other sites scoring over 100 and the cut sites are shown in column 4. **It should be noted that data for the acidic repeat region from 330-430 of LANA was excluded from the analysis due to the large number of DEED*DEED sites that score extremely high as caspase sites in SP. While these sites in LANA’s repeat region may be bona fide caspase sites, we did not find evidence for that in our previous study of LANA {Davis, 2015 #1520}.

| KSHV Protein | Top scoring site by Site Prediction | SP score for top site and caspase class (I, Ex, Inf) | Other SP sites scoring over 100 |
| --- | --- | --- | --- |
| ORF45* | DEED<sup>99</sup>*EDEE | 2787 (Ex) | D101, D106, D112, D248, D277 |
| K10.5/vIRF3 | EEVD<sup>88</sup>*DGAG | 2679 (Ex) | none |
| K5/MIR2* | DEPD<sup>222</sup>*GGPN | 1852 (I, Ex, Inf) | D37 |
| **ORF57/MTA\*** | **DETD<sup>33</sup>\*APTL** | **1742 (I, Ex, Inf)** | D307 |
| ORF38 | VDVD<sup>18</sup>*GEPL | 1649 (Ex) | none |
| ORF27* | VETD<sup>97</sup>*AI | 1146 (Ex) | D291 |
| ORF22* | DESD<sup>648</sup>*GLQS | 985 (I, EX, Inf) | D336 |
| ORF25* | VEGD<sup>345</sup>*KA | 926 (Ex) | D280 |
| ORF10 | VEGD<sup>208</sup>*PE | 863 (Ex) | none |
| ORF17* | VEKD<sup>518</sup>*APTP | 728 (Ex) | D366 |
| **ORF73 LANA\*** | **DSVD<sup>53</sup>\*GREC** | **687 (I, Ex)** | D324 ** |
| ORF63* | TEYD<sup>529</sup>*ED | 518 (Ex) | D5, D688 |
| ORF62 | VDLD<sup>208</sup>*ES | 504 (Ex) | none |
| ORF70 | VDAD<sup>174</sup>*AD | 487 (Ex) | none |
| ORF59 | ESPD<sup>332</sup>*SPPL | 464 (Inf) | none |
| ORF67 | VESD<sup>116</sup>*VY | 424 (Ex) | none |
| ORF37* | DTLD<sup>21</sup>*GLTV | 413 (Ex) | D341 |
| ORF64* | DESD<sup>1966</sup>*TASG | 392 (Ex) | D430, D998 |
| K8* | IEED<sup>28</sup>*LS | 320 (I, Ex) | D42 |
| ORF54 | GETD<sup>93</sup>*KD | 297 (Ex) | none |
| ORF50/RTA | DSPD<sup>408</sup>*NPSS | 276 (Inf) | none |
| K11/vIRF2* | LAPD<sup>27</sup>*SPRP | 274 (Inf) | D254 |
| ORF6* | TEED<sup>1039</sup>*VI | 262 (Ex) | D64, D400, D644 |
| ORF21/TK* | TDDD<sup>116</sup>*SG | 253 (Ex) | D45, D400 |
| ORF75* | IEDD<sup>184</sup>*VI | 252 (Ex) | D13 |
| ORF34 | EAVD<sup>139</sup>*GLCD | 226 (Ex) | none |
| ORF4 | SLTD<sup>495</sup>*SA | 217 (I, Ex, Inf) | none |
| ORF71/vFLIP | TDVD*ALMS | 214 (Ex) | none |
| K3/MIR1* | EDED<sup>5</sup>*VP | 183 (Ex) | D232 |
| ORF19 | LELD<sup>71</sup>*RL | 153 (Ex) | none |
| ORF20 | EVLD<sup>30</sup>*SSSE | 135 (Ex) | none |
| K14* | TDSD<sup>178</sup>*GLTV | 128 (Ex) | D266 |
| ORF74/GPCR | LDDD<sup>14</sup>*ES | 125 (Ex) | none |
| ORF2 | SSLD<sup>94</sup>*AALG | 115 (Ex) | none |
| ORF49 | QELD<sup>31</sup>*TL | 101 (Ex) | none |

### K5-FLAG undergoes caspase cleavage in αFas-treated BJAB-K5-FLAG cells

Because of its role in thwarting the immune response against KSHV, we were particularly interested in investigating the potential cleavage of K5/MIR2, an early lytic protein with the third highest SP score (DEPD^222^*GGPN) (Table 1). A primary function of K5 is to downregulate immune surface markers in KSHV-infected cells (15–18). To determine if the K5 protein might be susceptible to caspase cleavage in cells, we developed a K5-FLAG-expressing BJAB cell line with the 3XFLAG-tag (aa sequence: DYKDHDGDYKDHDIDYKDDDDK) at the C-terminus of the protein (Fig. S1). BJAB-K5-FLAG cells were treated with 10 ng/mL of αFas (a known apoptotic agent that works by activating initiator caspase-8 through the FADD pathway) in the absence or presence of a pan-caspase inhibitor (Z-VAD-FMK, called henceforth ZVAD) or caspase inhibitors directed at the three specific classes of caspases: C1, C3/C7, and C8. BJAB-K5-FLAG protein extracts were prepared and analyzed by immunoblot and probed using an anti-FLAG antibody (Fig. 1A). In untreated BJAB K5-FLAG cells, K5-FLAG was detected near the expected relative molecular weight of 32 kDa (expected MW is 31 kDa based on 28 kDa for K5 plus approximately 3 kDa for 3X FLAG C-terminal tail) (Fig. 1A, lane 1). When cells were treated with αFas for 24h, the levels of K5-FLAG protein decreased substantially (Fig. 1A, lane 2) and a new band was detected with the FLAG antibody at a relative molecular weight of about 6 kDa, indicative of caspase cleavage near the C-terminus of K5-FLAG (Fig. 1A, lane 2). In the presence of 25 µM ZVAD, the K5-FLAG levels were close to that of the untreated control and the 6 kDa band was no longer detected (Fig. 1A, lane 3), indicating that caspase cleavage of K5 had been blocked. A caspase-3/7 inhibitor decreased the production of the 6 kDa fragment, and a caspase-8 inhibitor it (Fig. 1A, lanes 5 and 6). By contrast, a caspase-1 inhibitor did not block αFas mediated production of the 6 kDa fragment (Fig. 1A, lane 4). Interestingly, while ZVAD and the caspase-8 inhibitor prevented the ⍺Fas-induced decrease in K5-FLAG and eliminated the production of the 6 kDa band, the caspase-3/7 inhibitor did not have this preventative effect, even though there were much lower levels of the 6 kDa band as compared to the αFas control (Fig. 1A, lane 5). This may indicate that cleavage is still occurring in the presence of the caspase-3/7 inhibitor but that subsequent degradation of the 6 kDa band also occurs under these circumstances.

**Fig. 1:**
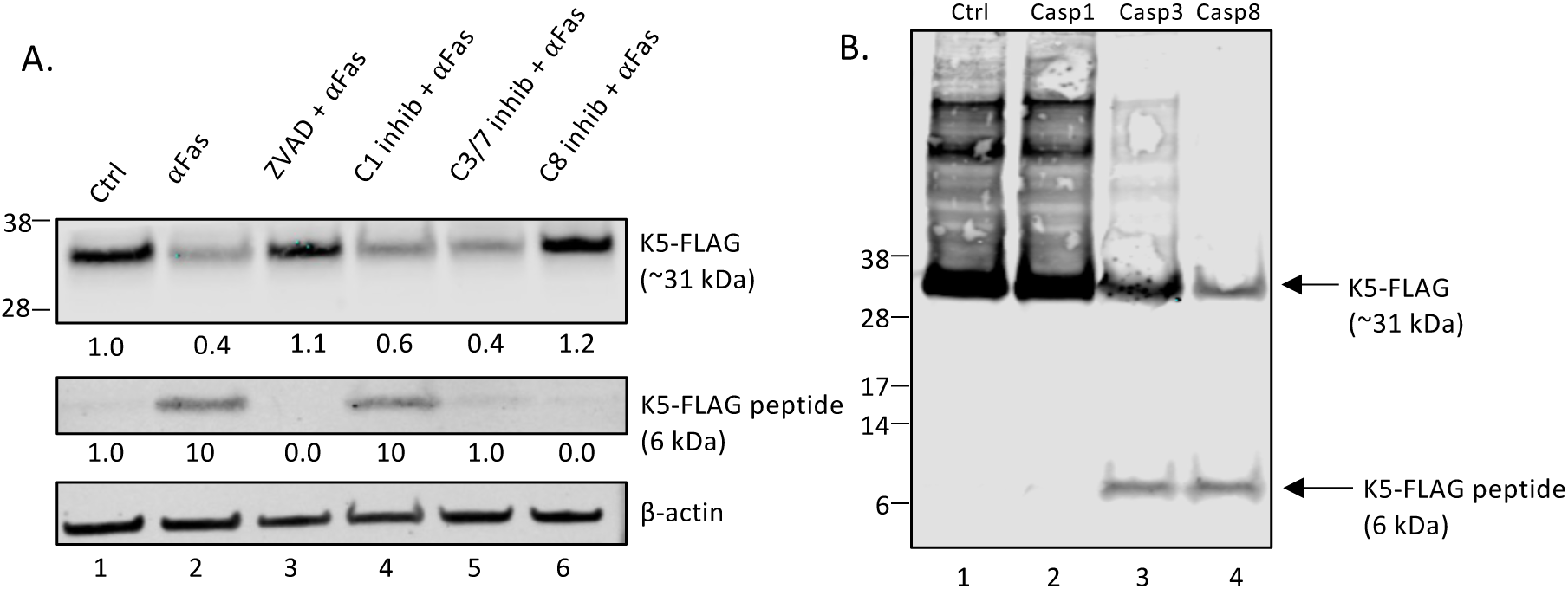
K5-FLAG undergoes caspase-cleavage in αFas treated BJAB-K5-FLAG cells and cell extracts. (A) Protein extracts were prepared from BJAB K5-FLAG cells (300,000 cells/ml) and analyzed by western blot following a 24h treatment with vehicle (DMSO) (lane 1), 10 ng/mL ⍺Fas (lane 2), 25 µM ZVAD for 2h followed by 10 ng/mL ⍺Fas (lane 3), 50 µM caspase-1 inhibitor followed by 10 ng/mL ⍺Fas (lane 4), 100 µM caspase-3/7 inhibitor followed by 10 ng/mL ⍺Fas (lane 5), or 25 µM caspase-8 inhibitor followed by 10 ng/mL ⍺Fas (lane 6). The blots were probed with anti-FLAG and β-actin antibodies. The signal intensity of the intact K5-FLAG band in lane 1 was normalized with respect to β-actin and the β-actin-normalized fold changes were calculated for the other treatments (B) Cytoplasmic protein extracts from BJAB K5-FLAG cells were prepared (without protease inhibitors) and then 8 µg of total protein in the presence of 50 mM DTT was either untreated or treated with 2 units of caspase-1 (lane 2), caspase-3 (lane 3), or caspase-8 (lane 4) for 1.5h and then stopped with SDS-sample buffer followed by analysis by western blot. The blot was probed with anti-FLAG antibody. The intact K5-FLAG and K5-FLAG peptide fragments are indicated.

To further investigate which class of caspases may be involved in the cleavage of K5-FLAG (initiator, executioner or inflammatory), we made cytoplasmic extracts of K5-FLAG in the absence of protease inhibitors and then tested the different classes of caspase for cleavage of K5-FLAG as assessed by western blot (Fig. 1B). K5-FLAG did not undergo cleavage by the inflammatory caspase-1. However, K5-FLAG was clearly processed by the executioner caspase-3 and the initiator caspase-8, and both generated a 6 kDa FLAG band like that seen in αFas treated BJAB-K5-FLAG cells. These results suggest that both executioner (caspase-3) and initiator (caspase-8) caspases can cleave purified K5-FLAG protein at the C-terminus. Interestingly, the highest SP scoring caspase site in K5 identified was at D222. Cleavage of K5-FLAG at D222 by caspases would generate an approximately 6 kDa K5-FLAG protein band like that observed by western blot (the calculated average mass for the C-terminal peptide with 3XFLAG is 6311) (Fig. S1). Also, both caspase-3 and caspase-8 scored high in SP for cleavage at D222 (scores of 1852 and 1690, respectively).

### K5-FLAG is Cleaved at D222 by Caspase-8

To identify the site of caspase cleavage in K5-FLAG, we prepared protein extracts from BJAB-K5-FLAG cells (Fig. 2A, lane 1). The extract was then treated with caspase-8 for 2 h to allow processing of K5-FLAG (Fig. 2A, lane 2). After caspase-8 processing, protease inhibitors (MG132, EDTA, and ZVAD) were added to prevent any further protease digestion that might otherwise occur in the extract. The sample was then dialyzed into Tris buffer and added to a slurry of anti-FLAG beads for 2 h at room temperature to bind any FLAG-containing peptide(s). The beads were washed 3 times with 0.1 M tris buffer and any FLAG-bound peptides and protein were eluted from the beads using 8 M urea acidified with 0.2% trifluoroacetic acid (TFA). The eluted material was then dialyzed into distilled water to remove the urea/TFA. The anti-FLAG western blot showed that this fraction contained a small amount of unprocessed K5-FLAG along with a 6kDa FLAG-containing peptide (Fig. 2A, lane 3). There was no peptide detected in the unbound fraction (Fig. 2A Lane 4). The intensity of the 6kDa band generated was always substantially lower than expected based on the intensity loss of the intact K5-FLAG after caspase-8 treatment. The reason for this is unclear but may be due to a relative inability of the anti-FLAG antibody to pick up the small peptide compared to that for larger proteins. However, it is also possible that subsequent degradation of the peptide occurs leading to a loss of detectable peptide.

**Fig. 2:**
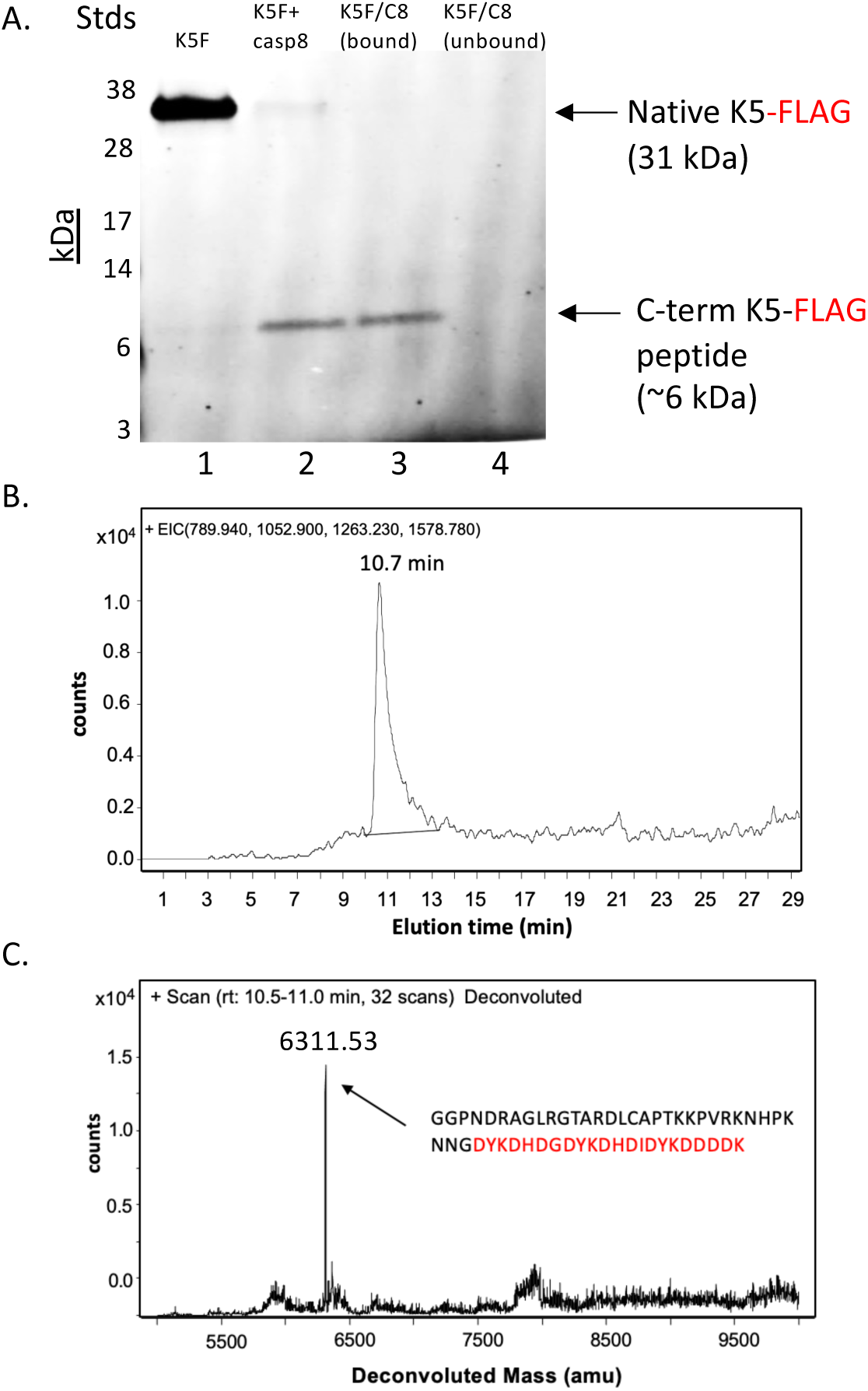
Purification of the K5-3XFLAG peptide and mass identification by RP/HPLC/MALDI TOF analysis. Protein extracts of BJAB-K5 cells were prepared using mPer and then used to purify K5-FLAG peptide using anti-FLAG beads. (A) Western blot for crude and purified K5-FLAG peptide using antiFLAG antibody. Protein extract of BJAB-K5 FLAG cells (lane 1), caspase-8 treated K5-FLAG mPer extract (lane 2), bound material eluted form anti-FLAG beads using 8M urea (lane 3), and the unbound flow thru fraction from anti-FLAG beads (lane 4). (B) Analysis of the purified K5-FLAG peptide by RP-HPLC/MS chromatography. The major ions predicted for the flag peptide produced following cleavage at D222 (789.8,902.5,1052.8,1263.1 and 1578.7) were extracted from the TIC chromatogram and a peak containing all the ions was identified eluting at 10.7 min. Major ions for the other predicted peptides produced if cut after D were not found. (C) Deconvolution of the peak identified in (B) revealed a molecular weight of 6311.53. Predicted molecular weight of 6310.70 for the peptide cleaved after D222 (amino acids 223-258).

The sample containing the FLAG-peptide was dried by speed-vac and then brought up in RP-HPLC running buffer (water with 0.1% formic acid and 0.02% TFA). The sample was then run on a C18 RP-HPLC column, and any bound peptides and proteins were eluted with an acetonitrile gradient and run directly into an Agilent MS/TOF mass spectrometer to identify masses for any eluting peptides. The TIC scan was interrogated for peptide ions consistent with cuts at aspartate residues in the C-terminus of K5-FLAG (Fig. S2). Only peptide ions consistent with a peptide generated from cleavage at D222 (789.8^8+^, 902.5^7+^,1052.8^6+^,1263.1^5+^, and 1578.7^4+^) were found (Fig. 2B) and deconvolution of the extracted ion peak gave a mass of 6311.51 (Fig. 2C), which is in very good agreement with the predicted average mass of 6310.70 for the D223-258-3xFLAG peptide. These data indicate that K5-FLAG is cleaved by caspase-8 at the highest scoring SP site (D222).

### Detection of the N-terminal cleavage fragment of caspase-cleaved K5-FLAG

We wanted to further confirm the cleavage of K5 through detection of the large N-terminal fragment that would be predicated to be generated following caspase-cleavage at the C-terminus. To this end, we utilized a rabbit antibody custom prepared by Genscript that targets amino acids 204-217 of K5, which is in the C-terminal region of K5 but prior to the D222 caspase-cleavage site. We first determined if this antibody was able to detect both K5 (as K5-FLAG) and the larger N-terminal fragment (amino acids 1-222) generated by caspase cleavage at D222 using extracts from BJAB and BJAB K5-FLAG expressing cells. Several non-specific bands were detected in BJAB and BJAB-K5-FLAG protein extracts using this rabbit antibody (Fig. 3A, lanes 1-3). However, the rabbit antibody clearly detected K5-FLAG in the BJAB-K5-FLAG protein extracts (Fig. 3A, lanes 4-6). Treatment with αFas led to a decrease in the total K5-FLAG detected and revealed a new band around 25 kDa indicative of the larger cleaved fragment (Fig. 3A, lane 5). When BJAB-K5-FLAG cells were first pretreated with the pan caspase inhibitor IDN-6556 before addition of αFas, then the loss of K5-FLAG was prevented and the 25 kDa band was no longer detected, and the levels of K5-FLAG were restored (Fig. 3A, lanes 5 and 6). The same blot was probed for K5-FLAG using an anti-K5 mouse monoclonal antibody (anti-K5 ms). The mouse monoclonal antibody was only able to detect K5-FLAG in the BJAB-K5-FLAG cells but not in BJAB cells (Fig. 3B, lanes 1 and 4). The mouse antibody, however, was not able to pick up the 25 kDa band or the 6 kDa FLAG-containing band upon treatment with αFas, even though it could detect a decrease in the amount of K5-FLAG suggesting the mouse antibody is only good at detecting intact K5-FLAG (Fig. 3B, lane 5). Although treatment with αFas led to a decrease in the amount of intact K5-FLAG detected using the mouse antibody, a new band was not detected, suggesting the mouse antibody can only detect intact K5-FLAG (Fig. 3B, lane 5). When cells were treated with the pan-caspase inhibitor IDN-6556 before addition of αFas, the loss of K5-FLAG, as determined using the mouse antibody, was prevented, again indicating caspase processing of K5 occurred following αFas treatment (Fig. 3B, lane 6). Together, these data indicate that K5-FLAG undergoes caspase cleavage near the C-terminus resulting in a 25 kDa truncated K5.

**Fig. 3:**
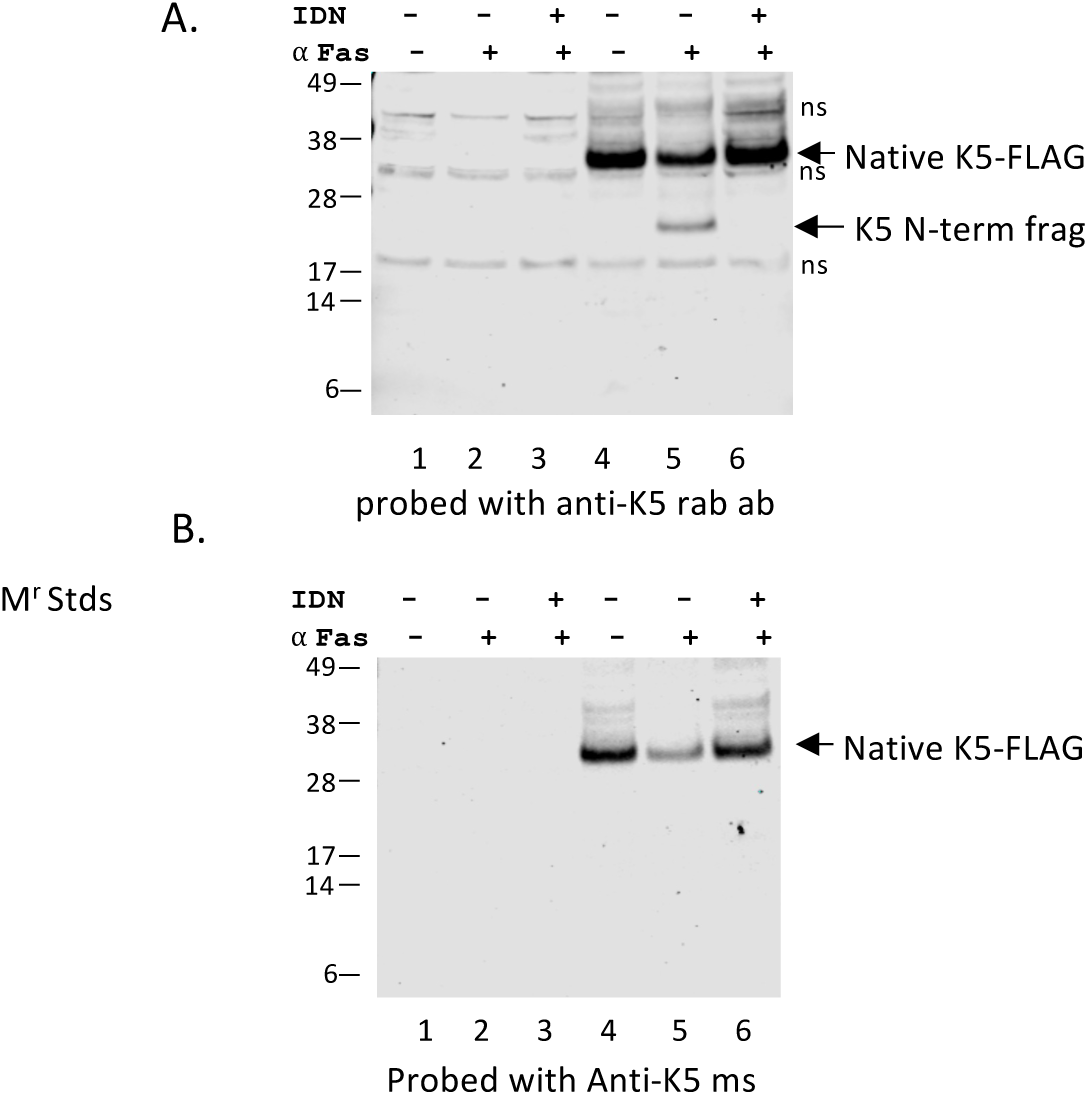
Analysis of BJAB and BJAB-K5-FLAG extracts by immunoblot using either a K5 mouse or K5 rabbit antibody. BJAB or BJAB-K5F cells were treated with vehicle controls, anti-Fas (10 ng/ml) or IDN-6556 pan-caspase inhibitor. Protein extracts were prepared 18h later and samples analyzed by immunoblot. (A) Blot probed with rabbit-anti-K5 antibody (anti-K5 rab ab) and (B) blot probed with mouse anti-K5 antibody (Anti-K5 ms). Lanes 1-3 are BJAB lysates and 4-6 are BJAB-K5F lysates. ns; non-specific band.

### KSHV K5 undergoes caspase cleavage in lytic PEL cells

Previous studies have established that lytic induction of KSHV-infected cells leads to caspase activation over levels that are seen in latency (9, 11–13). We therefore wanted to determine whether K5 undergoes caspase cleavage in KSHV-infected cells induced to lytic replication. To determine if K5 processing also takes place in the context of KSHV-infected PEL cells, we prepared cytoplasmic and nuclear extracts of BCBL-1 cells 48h after lytic induction with butyrate in the absence or presence of the pan-caspase inhibitor ZVAD, then analyzed for K5 with rabbit anti-K5 antibody (Genscript). While the mouse monoclonal antibody is useful for looking at the levels of intact K5, the rabbit antibody allows for detection of intact K5 as well as the large, cleaved form of K5. K5 levels increased 3.3-fold in cytoplasmic extracts upon induction with butyrate but also revealed a band running below intact K5 at around 25 kDa which is possibly cleaved K5 (Cl-K5?) (Fig. 4A, lanes 1 and 2). Treating BCBL-1 cells with ZVAD alone had no discernible effect on background K5 levels (Fig. 4A, lane 3). However, when BCBL-1 cells were pretreated with ZVAD followed by lytic induction with butyrate, full length K5 levels nearly doubled while the lower25 kDa band was less intense, indicating that a substantial portion of K5 produced during lytic activation undergoes caspase cleavage (Fig. 4A, lanes 2 and 4). We also examined the nuclear extracts of BCBL-1 cells (which consists of both nuclear and plasma membrane proteins) using the NE-Per kit from Thermo Fischer, since K5 is known to be shuttled to the inner plasma membrane from the endoplasmic reticulum (ER) (16) and has been found in the ER fraction of cells (19). K5 was also detected in the membrane fraction of butyrate-treated BCBL-1 cells and increased 5-fold over background levels from uninduced cells (Fig. 4B). K5 further increased to 7.5-fold over control in the presence of butyrate plus ZVAD while the levels of the lower band decreased. This again is consistent with caspase cleavage of K5 in the membrane protein extracts (Fig. 4B).

**Fig. 4:**
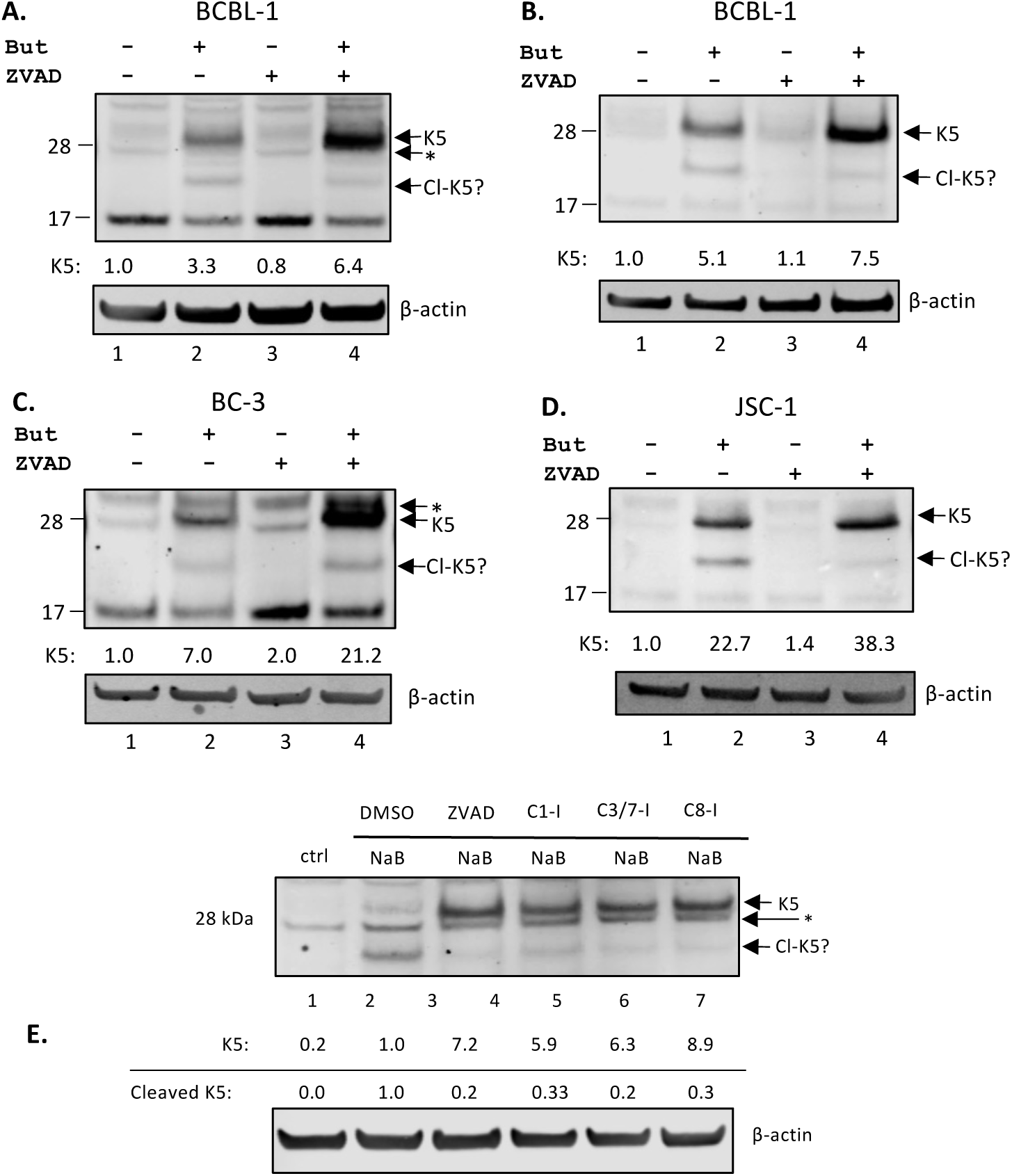
Caspase inhibition increases K5 protein levels in lytic PEL cells. (A) Cytoplasmic protein extracts and (B) nuclear protein extracts from BCBL-1 cells (C) cytoplasmic protein extracts from BC-3 cells and (D) Nuclear protein extracts from JSC-1 cells analyzed by western blot using the rabbit K5 antibody and β-actin antibody. Protein lysates were extracted from cells after 48h of treatment with DMSO control and PBS control (lane 1), DMSO and 1 mM butyrate (lane 2), 25 µM ZVAD and PBS (lane 3), or ZVAD and butyrate (lane 4). Signal intensity of each protein with respect to β-actin and normalized to lane 1 is displayed under each blot. (E) Cytoplasmic protein extracts from BCBL-1 cells analyzed by western blot using K5 rabbit antibody. Protein lysates were extracted from BCBL-1 cells after 48h of treatment with DMSO control and PBS control (lane 1), DMSO and 1 mM butyrate (lane 2), 25 µM ZVAD and butyrate (lane 3), 50 µM caspase-1 inhibitor and butyrate (lane 4), 100 µM caspase-3/7 inhibitor and butyrate (lane 5), or 25 µM caspase-8 inhibitor and butyrate (lane 6). Signal intensity of each protein in lane 2 was normalized with respect to β-actin and set as 1.0 and then the fold differences calculated for the different treatments and is displayed under each blot. The putative cleaved K5 band is labeled as CL-K5 and runs at approximately 25 kDa. Non-specific bands detected with the rabbit anti-K5 ab are indicated with an (*).

In BC-3 cells, K5 levels increased 7-fold with butyrate induction but in the presence of ZVAD, K5 increased more than 21-fold over control or three times that seen with butyrate alone (Fig. 4C). By contrast to BCBL-1 cells, the lower band seen in butyrate-treated BC-3 cells increased in intensity in the presence of ZVAD although the ratio of the lower band to the upper band was measurably decreased (Fig. 4C, lanes 2 and 4). We also examined JSC-1 cells, which are co-infected with EBV and KSHV (20). We only show the nuclear extracts for JSC-1 cells since the cytoplasmic extracts yielded many nonspecific bands around the K5 region, making it more difficult to quantitate K5 (although caspase processing of K5 was observed) (Fig. S3). Butyrate strongly induced K5 (almost 23-fold) in JSC-1 nuclear extracts when compared to the uninduced control (Fig. 4D, lane 2), and in the presence of ZVAD, the K5 levels increased to 38-fold over the control without butyrate (and 1.4 -fold over butyrate alone). As with BCBL-1 cells, the lower K5 band decreased in the presence of ZVAD which is consistent with it being a caspase-cleaved form of K5 (Fig. 4D, lane 4). Note that caspase cleavage at D222 in K5 would result in a small C-terminal peptide (3 kDa) and a larger truncated form of K5 with a molecular weight of about 25 kDa, which is close to the molecular weight of this unidentified lower band observed with the rabbit antibody (estimated 25 kDa based on M_r_ markers) for this unidentified lower band in these samples. These data provide good evidence that K5 undergoes processing near the C-terminus by caspases in PEL cells during lytic replication.

Further experiments were done to explore which caspases may be involved in cleavage of K5 during lytic replication by using inhibitors for the specific caspases. BCBL-1 cells were treated with butyrate without and with specific caspase inhibitors representing each of the three classes of caspases: inflammatory (capsase-1), apoptotic initiator (caspase-8), and apoptotic executioner (caspases-3/7). These cell extracts were analyzed by western blot and probed for K5 expression using rabbit anti-K5 antibody (Fig. 4E). In control BCBL-1 cells, there was low K5 expression, as seen previously. Following lytic induction with butyrate, we detected some K5 expression at the expected relative molecular weight of 28kDa, but more protein was detected at the 25 kDa band. Again, ZVAD increased full-length K5 and decreased the levels of the 25 kDa band. Comparison of the three different caspase inhibitors used at their recommended optimal concentrations demonstrated that inhibitors of caspases-3/7 and caspase-8 were the most effective at blocking cleavage and increasing native K5 protein (Fig. 4E). As we saw with BJAB-K5-FLAG extracts (Fig. 1) the caspase-1 inhibitor also had some activity but was the least effective of the caspase inhibitors as there was less native K5 and more of the 25 kDa fragment than with the other three inhibitors (Fig. 4E). Overall, these data with the different caspase inhibitors suggests there is some caspase cleavage at the C-terminal site by caspases from all three classes of caspases. However, the accumulation of native K5 and substantial reduction of the 25 kDa band by caspase-3/7 and −8 inhibitors provides evidence that the apoptotic executioner caspases-3/7 and initiator caspase-8 are likely to be primarily responsible for the cleavage of K5 in lytic PEL cells.

### BJAB cells expressing K5-FLAG are resistant to caspase-mediated αFas induced cell death

The experiments with K5-FLAG indicated that K5 could be efficiently cleaved by caspase-3 and caspase-8. Caspase-8, an initiator caspase that activates downstream effector caspases such as caspase-3 and 7, can be activated by treatment of cells with αFas (21). We examined whether expression of K5-FLAG in BJAB cells may affect caspase-mediated cell death caused by αFas. Treatment of BJAB cells for 24h with αFas led to significant cell death (>50%) as measured by trypan blue exclusion and this was significantly inhibited with the pan-caspase inhibitor ZVAD, indicating that the death occurring was caspase-mediated (Fig. 5A). BJAB cells expressing K5-FLAG also underwent significantly less cell death (<10% compared to its control) when treated with αFas suggesting K5-FLAG delays or prevents caspase-mediated cell death (Fig. 5A). Similar results were obtained when measuring αFas-induced cell death by FACS analysis using PI staining (Fig. 5B).

**Fig. 5:**
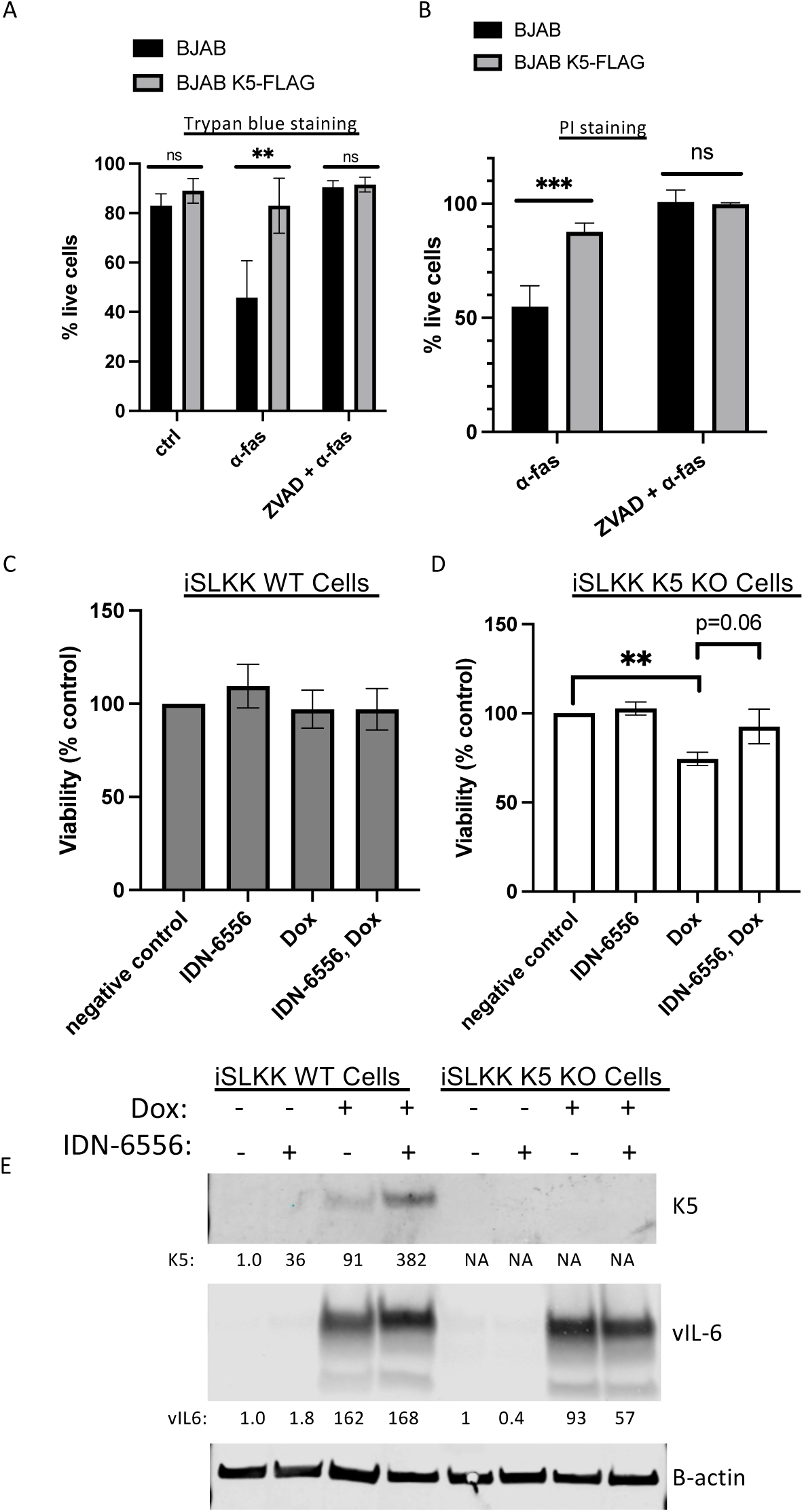
BJAB cells expressing K5-FLAG are resistant to caspase-mediated αFas induced cell death while K5 KO cells are susceptible to caspase-mediated cell death in cells induced to lytic replication by doxycycline. BJAB and BJAB-K5FLAG cells were plated at 200,000 cells per ml and treated with PBS control, αFas (10 ng/ml), or αFas in the presence of 25 µM ZVAD. After 24h the cell viability was assessed by (A), trypan blue staining or (B) PI staining by FACS analysis. (C) Bac16 iSLKK wildtype (WT) and (D) K5 knockout (KO) cells were pretreated for 2h with DMSO or IDN-6556, then treated with diluted DMSO or 1µg/mL doxycycline (Dox). After 48h, floating cells were collected, and adherent cells were removed with trypsin and collected. The combined solutions of floating and adherent cells were counted using trypan blue staining and cell viability was calculated. Shown are the averages and standard deviations for 3 separate experiments ** P< 0.01, *** P< 0.005. (E) Western blot for K5 (using the mouse antibody) and vIl-6 (used as another a lytic marker) in WT and K5-KO cells 48h after treatment with Dox. The relative values for K5 and vIL-6 are indicated and are normalized to their respective controls.

### delta-K5 iSLKK cells are susceptible to caspase-mediated cell death in cells induced to lytic replication by doxycycline

Following up on this, we were interested in knowing if K5, produced during lytic replication, could play a similar role in protecting KSHV-infected cells from caspase-mediated cell death. To address this question, we obtained WT and K5-knockout KSHV-infected iSLK cells (a kind gift from Dr. Jae Jung (22)). We first examined the level of expression of K5 transcripts in WT cells and in the K5-knockout cells. We followed the change in K5 gene expression following induction of the cells with Dox which leads to activation of an RTA-expressing plasmid in these cells. In WT cells, K5 transcripts on day 0 were about 10^4.2^ copies per µg of RNA and the levels steadily increased to over 10^7^ copies per µg by 72 hours (Fig. S4A). However, no transcripts were detected in the K5-knockout cell line confirming the knockout line did not express K5 mRNA (Fig. S4A). In addition, we examined these cells for K5 protein expression following induction with Dox using the rabbit and mouse K5 antibodies. Using the rabbit antibody to K5, K5 was only detected in the WT induced cells but not in the K5 knockout line although the antibody picked up a strong non-specific band just below K5 that was seen in all samples (Fig. S4B) Using the mouse antibody a single band for K5 was detected only in the Dox-induced iSLKK WT cells but not in the K5 knockout cells (Fig. S4C). We did not observe a truncated/caspase-cleaved form of K5 in the WT infected cell extracts using the rabbit anti-K5 antibody but the low levels of K5 induced in this system may preclude its detection. In addition, the rabbit antibody reacted to a nonspecific band just below K5 possibly obscuring detection of the cleaved form (Figure S4B). We also examined the extracts for the lytic ORF45 tegument protein. ORF 45 was detected in WT and K5-knockout iSLK cells at similar levels suggesting similar lytic activation in both cell lines following Dox treatment (Fig. S4D).

Previous studies have demonstrated that induction of lytic replication by Dox in iSLKK cells does not lead to cell death even though caspase activation is observed (12). We wanted to determine if low dose Dox (1 µg/ml) might induce cell death in cells lacking K5. To explore this, we treated the two lines with Dox (1 µg/ml) without or with the pan caspase inhibitor IDN-6556 (10 µM) which has been shown to inhibit caspase activity in the WT iSLKK lines (12). WT iSLKK infected cells did not undergo significant cell death following lytic activation with Dox for 48 h, (Fig. 5C). However, the K5-knockout cells underwent significant (average 25%) cell death in this period as compared to the control cells (Fig. 5D). This Dox-induced cell death was mostly blocked in the presence of IDN-6556 and the cell death in the IDN-6556/Dox treatment was no longer significantly different than the uninduced control (Fig. 5D). These data suggest that K5 in iSLK cells infected with WT KSHV can at least, in part, prevent caspase-mediated cell death during lytic replication. We also measured the levels of K5 in WT and K5 knockout cells following Dox treatment for 48h. Using our mouse K5 antibody, K5 was detected in Dox-treated WT cells but not in K5 knockout cells (Fig. 5E). K5 levels in iSLKK WT cells increased more than 3-fold in the presence of IDN-6556 implying caspase cleavage of K5 was occurring in Dox treated cells. We also looked at the levels of vIL-6, another lytic protein. By contrast to K5, the levels of vIL-6 did not substantially increase in the presence of IDN-6556 in WT or K5 knockout cells (Fig. 5E). Together, these data indicate the presence of K5 can delay or prevent caspase-mediated cell death in cells undergoing lytic replication and that the absence of K5 can result in caspase-mediated cell death in these cells.

### K5 may reduce caspase-mediated cell death by delaying caspase cleavage of downstream caspase targets

Since we found that the presence of K5-FLAG decreased αFas induced cell death, we measured the levels of caspase-8, and caspase-3 and the cleavage of their downstream targets such as PARP and caspase-6 in BJAB and BJAB-K5-FLAG cells. After 4h of αFas stimulation, we detected similar caspase-8 activation based on western blot analysis which revealed the activation doublet 43/41 (Fig. 6A), suggesting K5-FLAG was not interfering with the initial activation of caspase-8 through the FADD receptor. Within the same samples, we also measure the activation of caspase-3 after 4h which occurs via caspase-8 cleavage of procaspase-3 during αFas treatment. Here, the induction of cleaved caspase-3 was somewhat lower (23.6-fold vs 18.6-fold) in the BJAB-K5-FLAG cells (Fig. 6B). We also examined the extent of PARP cleavage in αFas treated BJAB and BJAB-K5-FLAG cells. PARP is cleaved by executioner caspases (3 and 7) and is a useful readout for caspase-mediated apoptosis (23, 24). The increase in cleaved PARP following αFas induction in BJAB K5-FLAG cells was less than one third of that seen in BJAB cells (2.0 vs 6.8), suggesting that K5 may in fact interfere with downstream targets of caspases (Fig. 6C). We also assessed the activation of caspase-6, which occurs via caspase-3 cleavage of procaspase-6. We were not able to detect cleaved forms of caspase-6, so we analyzed the loss of uncleaved procaspase-6 as an indirect measure. At 4h post αFas treatment, we detected similar decreases in procaspase-6. (Fig 6D). We also looked at the levels of procaspase-6 after 18h of treatment with ⍺Fas. At this time point, we did see a somewhat greater loss of procaspase-6 in BJAB cells as compared to the BJAB-K5F cells suggesting greater processing in BJAB cells to active caspase-6 (Fig.6E).The loss of procaspase-6 was prevented when αFas treatment was done in the presence of the pan caspase-inhibitor IDN-6556 indicating a caspase-dependent conversion of procaspase-6 to its active forms (Fig.6E). Taken together, these data suggest that K5FLAG may delay caspase-3/7-mediated events and thus delay caspase-dependent cell death.

**Fig. 6:**
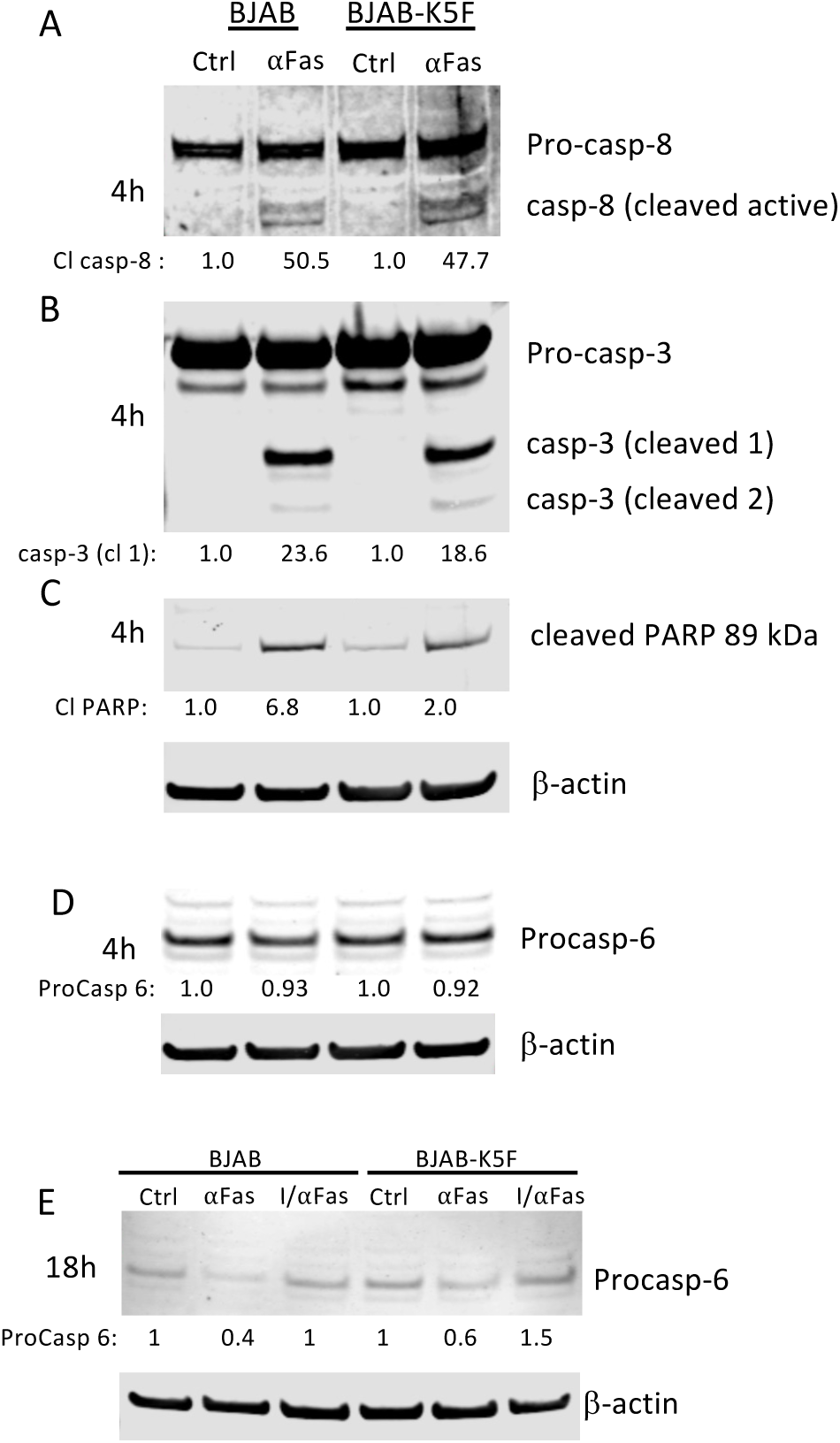
Downstream targets of caspase-3/7 are cleaved less in BJAB K5F cells. RIPA protein extracts were prepared 4h or 18h after treatment of BJAB cells or BJAB K5F cells with ⍺Fas (10 ng/ml) and run on western blots and analyzed for, (A) caspase-8 and cleaved caspase-8, (B) Caspase-3 and cleaved caspase-3, (C) the cleaved form of PARP (89 kDa) and (D) procaspase 6. Signal intensity of each protein of interest was normalized with respect to β-actin and the value is displayed under each blot. Signal intensity of each protein with respect to β-actin and normalized to lane 1 is displayed under each blot. (E) An 18h experiment was done with ⍺Fas treatment without or with IDN-6556 (10 µM) caspase inhibitor (labeled I/⍺Fas) and then analyzed for procaspase-6. The relative levels of procaspase-6 are shown after normalizing to β-actin.

### Truncated GFP-K5 still can downregulate the immune surface markers MHC-1, ICAM-1, and B7-2

Based on our data, the caspase-cleavage site identified in K5 yields a large N-terminal fragment of 25 kDa (predicted MW of 24.4 kDa) and a small peptide of about 3.6 kDa (or 6 kDa in case of K5-FLAG). We decided to explore if loss of the C-terminal peptide due to caspase cleavage affects the ability of K5 to down-regulate immune surface markers, which is considered the primary function of K5 (15–17, 22, 25). We developed plasmids expressing GFP-K5 or a truncated form of GFP-K5 representing the caspase-cleaved large fragment designated as GFP-K5-D222 and then measured by FACS analysis their ability to downregulate MHC-I in HEK-293 cells. GFP-K5 and GFP-K5 D222 were both well expressed (Fig. 7A). Also, both were able to significantly downregulate MHC-I (Fig. 7B). Although we explored the effect of GFP-K5’s on the surface markers ICAM-1 and B7-2 in these cells, the levels of expression of these surface markers in control cells were too low to discern clear differences between the GFP-K5’s activity. We also evaluated the effect of the GFP-K5s on MHC-I, ICAM-1 and B7-2 expression in BJAB cells, since these cells have higher expression of ICAM-1 and B7-2. In this case, as with HEK-293 cells, both GFP-K5 and GFP-K5-D222 downregulated MHC-I to similar extents. Moreover, they were equally effective at down-regulating ICAM-1 and B7-2 in BJAB cells (Fig. S5). We also explored whether blocking caspase processing of K5, in the context of virus infected cells, might affect the extent of downregulation of MHC-I in BCBL-1 cells during lytic replication. In two separate experiments, pretreatment of the cells with ZVAD had little effect on the downregulation of MHC-I in lytic BCBL-1 cells, going from an average of 63% downregulation by butyrate alone to 71% downregulation in the presence of ZVAD and butyrate (Fig. 7C). Overall, these data suggest that caspase cleavage of K5 does not substantially impair its ability to down-regulate immune surface markers and that the main advantage of this cleavage for KSHV may be to thwart caspase-mediated cell death during lytic replication.

**Fig. 7:**
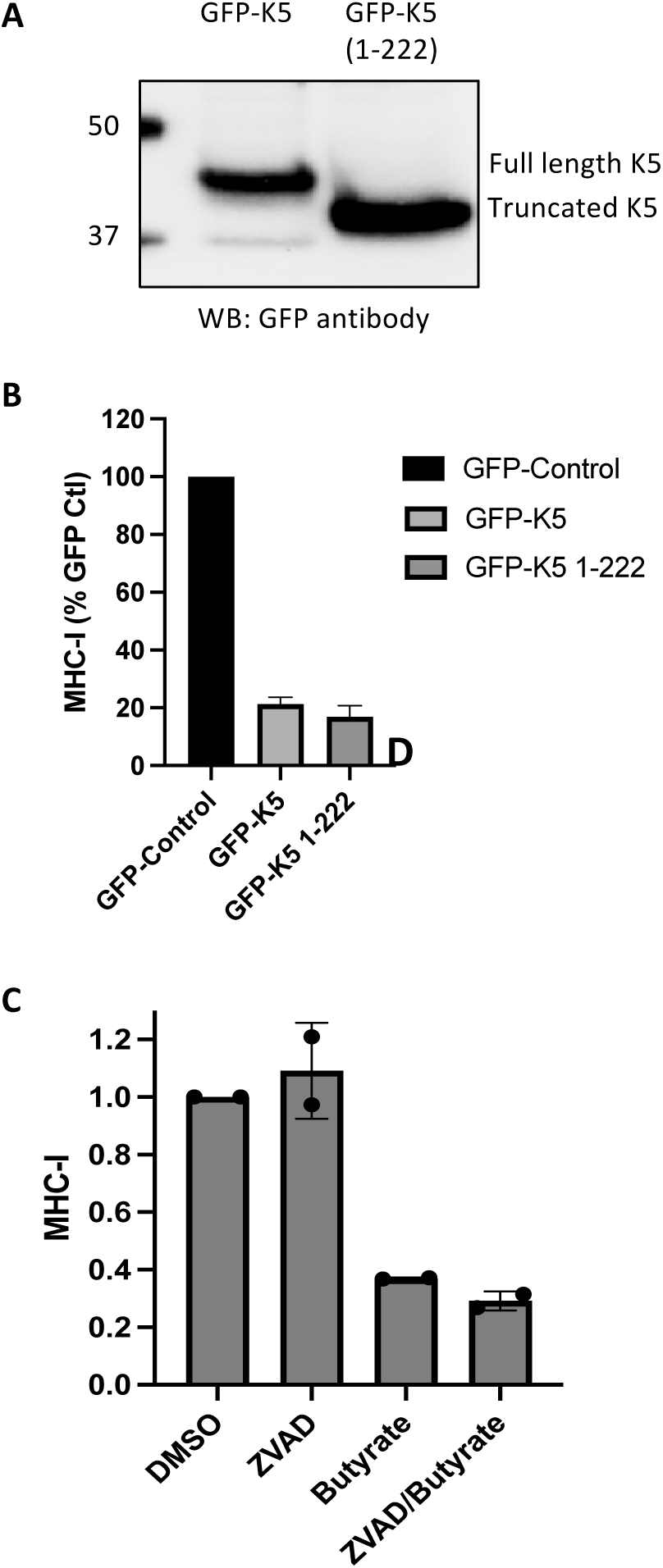
GFP-K5 and truncated GFP-K5 induce the downregulation of MHC-I. (A) Immunoblot for GFP-K5 and truncated GFP-K5 detected by using GFP antibody. (B) MHC-I surface expression in HEK-293T cells measured by FACS analysis 28h after transfection with GFP-control vector, GFP-K5 plasmid, or truncated GFP-K5 (GFP-1-222) plasmid. (C) MHC-I surface expression in BCBL-I cells measured by FACS analysis following treatment for 48h with DMSO control, ZVAD (25 µM), butyrate (1 mM), or ZVAD then butyrate. ZVAD was added 2h prior to treatment with butyrate. The data plotted in B is the average and standard deviation from 3 separate experiments while in C the data is the average of two experiments with each value shown as a dot.

## Discussion

One of the most studied mechanisms of cellular defense against viral pathogens is caspase-mediated cell death where cells either undergo apoptosis, pyroptosis, or necroptosis upon infection. To survive infection, viruses must find ways to prevent these forms of caspase-mediated cell death to facilitate their replication and survival. Several nice reviews have been published that describe the variety of ways in which human and non-human viruses evade caspase-mediated cell death (1, 3, 4, 26). The most direct way that viruses do this is by encoding viral proteins that directly inhibit caspases as is the case the baculovirus-encoded P35 (reviewed in (1)). However, this is not the only way caspases are thwarted. In some cases, cleavage of viral proteins by caspases has been shown to yield cleavage products with novel activities. For some viruses, (AMDV, HPV, SARS-CoV etc.) caspase cleavage products have been shown to enhance viral replication, for example by enhancing nuclear translocation of viral proteins (reviewed in (3)). In some cases, cleavage of viral proteins by caspases has been shown to lead to products that attenuate viral replication (13, 27, 28).

Previous studies by our group and others showed that KSHV encodes at least two proteins, LANA and ORF57, that are cleaved by caspases. Cleavage of LANA was shown to inhibit caspase activity, while evidence was provided that cleavage of ORF57 could yield a cleavage product that reduces lytic replication (13). In this study, we used SP (https://www.irc.ugent.be/prx/bioit2-public/SitePrediction/index.php) to identify additional KSHV-encoded proteins that may undergo caspase cleavage. Using SP, the top ten KSHV proteins predicted to be susceptible to caspase cleavage were ORF45, K10.5, K5, ORF57, ORF38, ORF27, ORF22, ORF25, ORF10 and ORF17; these all scored higher than the confirmed N-terminal caspase-3 cleavage site in LANA (D53). In this work, we confirmed that K5 is indeed cleaved by caspases in PEL cells undergoing lytic replication. K5 is an early lytic KSHV protein that, along with K3, down-regulates immune surface markers during lytic replication(15–18, 25). Utilizing K5-FLAG protein expressed in lentivirus-infected BJAB cells, we showed that K5 is cleaved at D222 by caspase-8 and caspase-3 but not caspase-1. Expression of K5-FLAG significantly decreased the extent of caspase-mediated cell death induced by αFas, an activator of caspase-8, and αFas treatment led to significant caspase-mediated processing of K5-FLAG. In KSHV-infected iSLK cells exposed to Dox, knock-out of K5 led to caspase-mediated cell death that does not occur in WT KSHV-infected cells exposed to similar concentrations of Dox (11). Our data suggest that K5 may contribute to the prevention of cell death during lytic replication of KSHV.

We originally became interested in K5 when we found that the immune modulator, pomalidomide, could prevent the downregulation of immune surface markers caused by K5 (29). We initially postulated that following their activation by pomalidomide, caspases may cleave and inactivate K5, resulting in increases in immune surface markers. However, we found that even after truncation at the D222 caspase cleavage site, K5 was able to efficiently downregulate immune surface markers. Consistent with this, others have shown that deletion of the C-terminal tail (amino acids 233-256) of K5 does not significantly alter its ability to downregulate MHC-I, ICAM-1 or IFN-gamma receptor 1(25). We also found that blocking caspase activity in KSHV-infected cells undergoing lytic replication did not significantly change the downregulation of surface markers. These data suggest that caspase cleavage of K5 at the C-terminus does not significantly impair its ability to downregulate immune surface markers.

While caspase-mediated cell death is typically associated with apoptosis (extrinsic and intrinsic), it is now appreciated that caspases also play a role in other types of cell death including pyroptosis, necroptosis and necrosis (30) (31–36). Given that caspase cleavage of K5 does not reduce its ability to downregulate immune surface markers but reduces caspase activity, we hypothesize that KSHV has evolved to encode caspase-cleavage sites as a mechanism to mute the overall caspase response, therefore blocking the cellular anti-viral strategy of caspase-mediated cell.

Interestingly, KSHV encodes for a variety of different caspase cleavage sites that are targeted by different caspase classes, and extending the observations here, one can postulate that this is a general mechanism for KSHV to divert caspases away from their usual functions which can thwart viral replication and persistence. Although we did not fully explore which types of caspase-mediated cell death were inhibited by K5, we did learn that it can prevent αFas-mediated cell death which usually goes through the extrinsic pathway of apoptosis by activating caspase-8 followed by subsequent activation of extrinsic caspases (caspase-3,6,7) and cleavage of PARP (23, 24). The increase in the amount of cleaved PARP was greater in control cells than in K5-expressing cells, although K5-expressing cells had higher baseline levels of cellular cleaved PARP to begin with. We also saw a delay in procaspase-6 processing in K5-FLAG-expressing cells. Caspase-6, which degrades Lamin A during apoptosis, can be directly activated by caspase-3 (37), and in K5-FLAG expressing cells the extent of caspase-6 activation also appeared to be impaired. Our data suggest that K5 doesn’t substantially impair caspase-8 or capsase-3 activation by αFas in cells but instead appears to decrease the cleavage of downstream cellular targets of caspase-3, like PARP and caspase-6, perhaps by acting as a decoy substrate. The evidence points to K5 as acting as a decoy substrate for caspase-3 although other mechanisms may be in play and require further investigation.

Although we demonstrate a role for K5 in preventing caspase-mediated cell death, it remains possible that the cleaved forms of K5 or other KSHV proteins play other important roles in viral replication, possibly by conferring new functions to these proteins. Previous work by others has described a role for caspase activation in enhancing KSHV infection by preventing type 1 interferon responses (11, 12). It is possible that one or more of the other KSHV proteins susceptible to caspase cleavage may play a role in this response to caspase activation. Both KSHV encoded K11/vIRF2 and K10.5/vIRF3 have high SP scores for predicted caspase cleavage sites (Table 1) and perhaps caspase cleavage of these proteins may play a role in altering the interferon response.

Interestingly, K10.5/vIRF-3, which had the second highest SP score at D88 (score 2679) (Table 1) is a latent protein that is only expressed in KSHV-infected B-cells like those seen in PEL and MCD (38). K10.5/vIRF-3 has already been shown to antagonize apoptosis by inhibiting p53 and has been shown to decrease caspase-3 activation (39, 40). The presence of a caspase cleavage site in vIRF-3 and vIRF-2, if functional, could provide additional mechanisms by which these viral proteins may disrupt or redirect caspase-mediated events and is ripe for future studies.

In summary, in this study, we show that KSHV K5 can be cleaved by caspases and that by reducing caspase activity, this cleavage provides an additional mechanism by which KSHV thwarts cellular innate immunity. We also describe predicted caspase cleavage sites in many other KSHV proteins. Further analysis of these interactions may yield other insights on the complex interplay between KSHV and host cells.

## Acknowledgements

This work was supported by the Intramural Research Program of the National Institutes of health, National Cancer Institute. We also thank Dr. Hong Seok Choi for his help in generating the BJAB-K5-FLAG cells.

## Materials and Methods

### Identifying potential caspase-cleavage sites in KSHV proteins

SitePrediction (SP) (14) is a web-based program to predict caspase cleavage sites. SP predicts cleavage sites for caspases 1, 3, 6, 7, and 8. As of this writing (January, 2025), SP is still available for public use at https://www.irc.ugent.be/prx/bioit2-public/SitePrediction/. However, previous programs like Cascleave, Cascleave-2, and others are no longer freely accessible online. Eighty-seven different KSHV encoded proteins were analyzed by SP and the prediction scores ranked from highest to lowest.

### Caspases and caspase inhibitors

Recombinant human caspase-1 (Cat# CC126), caspase-3 (Cat# CC119), and caspase-8 (Cat# CC123), as representatives of the three different classes of caspases, were from Sigma Aldrich-Calbiochem (St Lousi. MO). Caspase-1 inhibitor (cat# 400015), caspase-3/7 inhibitor (Cat# 218832), caspase-8 inhibitor (cat# 218759), cell permeable pan-caspase inhibitor ZVAD-FMK (cat# 627610) and αFas (Cat# 05-201) were from EMD-Millipore Sigma. The pan-caspase inhibitor IDN-6556 (Cat# S7775) was from Selleck Chemicals. All inhibitors were dissolved into 100% cell-culture grade DMSO and stored at −20°C.

### Cells, cell culture, and reagents

BJAB, BC-3 and JSC-1 cells were from the American Type Culture Collection (ATCC), Manassas, VA). BCBL-1 cells were obtained from the NIAID AIDS Research and Reagent Program (Rockville, MD). Cells were cultured in RPMI 1640 medium (Life Technologies, Carlsbad, CA) with 15% heat-inactivated fetal calf serum (HyClone,Waltham, MA), 1% penicillin streptomycin glutamine (1000 units/ml penicillin, 10000 µg ml^-1^ streptomycin, 29.2 mg/ml L-glutamine) (Life Technologies). BJAB cells constitutively expressing K5-FLAG cells were prepared using lentivirus encoding a puromycin resistance marker as well as full length K5 containing a 3XFLAG C-terminal tag (9332 bp total) from Sigma. Lentivirus was electroporated into BJAB cells and cells expressing K5-FLAG, driven by EF1A promoter, were selected with puromycin. Confirmation of K5-FLAG expression was done using an antibody to 3XFLAG (cat# F1804) from Sigma as well as a mouse monoclonal antibody toward K5 (328C7) which was obtained from Drs Koichi Yamanishi and Keiji Ueda (Osaka University Medical School, Japan) (19). WT and K5-knockout infected iSLK cells were obtained as a kind gift from Dr. Jae Jungs group (22). Quantitative PCR was carried out described previously and the primers used for K5, and the controls are listed (Fig S5). A rabbit K5 antibody directed toward a K5 peptide, amino acids 206-218, was generated by Genscript (Piscatawy, NJ). A monoclonal β-actin antibody (cat# A5441) and sodium butyrate (NaB), which was used to induce lytic replication in KSHV-infected cells, were from Sigma-Aldrich (St Lousi, MO). An antibody directed toward ORF45 was from Abcam (Cambridge, MA, Cat.# ab36618). Antibodies to caspase-8 (cat# 9496, rab), caspase-7 (cat# 9492, rab), full-length caspase-6 (cat# 9762, rab), cleaved capsase-6 (cat# 9761, rab), caspase-3 (cat# 14220, rab), PARP (cat# 9532, rab), and cleaved PARP (cat# 9741, rab), were from Cell Signaling (Boston, MA). A monoclonal antibody to KSHV ORF2/vIL-6 was prepared as described previously (41). A synthetic peptide encompassing amino acids 223-278 of K5-FLAG was prepared by Vivitide (Gardner, MA).

### Purification of K5-FLAG peptide following caspase cleavage

K5-FLAG underwent caspase cleavage by caspases-3 and -8 but not caspase-1. To identify the site of caspase cleavage, the approximately 6 kDa FLAG containing peptide generated following treatment with caspase-8 and detected by western blot was purified from Anti-Flag labeled magnetic beads form Sigma (Cat.# M8823). Protein extracts of K5-FLAG expressing cells were prepared using mPer protein extraction reagent from Pierce and done in the absence of protease inhibitors. The mPer extracts were then treated with caspase-8 and the generated FLAG-peptide was captured using the anti-FLAG beads. The bound peptide was eluted with 8 M urea and then dialyzed against distilled water to remove the urea. The solution was then dried by speed vac and put up into RP-HPLC/MS running buffer (0.1% formic acid and 0.02% trifluoroacetic acid). The solution was injected onto a VyDac C18 column (Cat# 218TP5205) and eluted with an acetonitrile (ACN) gradient (containing 0.1% formic acid and 0.02% trifluoroacetic acid). Analysis was done for 5 min at 0% ACN followed by a 2.5%/min increase for the next 30 minutes. Peptides were analyzed in-line with an Agilent MALDI-TOF mass spectrometer and were then identified by extracting the predicted molecular ions for peptides generated from the K5-FLAG C-terminus following caspase cleavage after the various aspartic acid residues (Fig. S2).

### Cell death and FACS analysis

To induce extrinsic apoptosis through the FADD pathway, we treated BJAB and BJAB-K5FLAG cells (200,000 cells/ml) with 10 ng/ml of αFas antibody in the presence and absence of 25 µM ZVAD-FMK for 24h Cell viability was then determined using trypan blue (Life Technologies, Grand Island, NY) staining as well as PI staining by FACS analysis as described previously (42). Cell death that was prevented with ZVAD-FMK was considered caspase-mediated cell death.

### Immunoblotting

Nuclear and cytoplasmic extracts as well as mPER extracts were prepared as described previously (9) and were made in the presence of protease inhibitors (Halt Protease Inhibitors Cocktail Kit, Pierce) and 5 mM EDTA unless otherwise indicated. Protein concentrations were determined using the BCA assay (Pierce) and samples were separated on 4-12% NuPAGE gels and transferred to nitrocellulose membranes by iBlot (Life technologies). Membranes were probed with antibodies directed toward β-actin, 3XFLAG, K5, K3, and ORF45 as indicated in the text. Blots were then incubated with appropriate secondary antibodies conjugated to alkaline phosphatase and visualized using stabilized Western Blue substrate (Promega) or with a goat anti-mouse or rabbit IR700 or IR800 secondary antibody (diluted 1:10,000) as indicated. Membranes exposed to LI-COR secondary antibodies were scanned using a LI-COR Odyssey infrared scanner. Images were analyzed using Image Studio 2.1 software.

### Transfection of HEK-293 cells

HEK-293 cells were plated at 12×10^5^ cells per well in a 6-well plate. After 24 hours, the cells were transfected using Genjet reagent (Cat# SL100488) from SignaGen (Frederick, MD) as per the manufacturer’s protocol. Briefly, DNA (Ctrl, WT-K5, or M-K5 expressing plasmids) and Genjet reagent were diluted separately by adding 2 µg DNA or 6 µL Genjet reagent to 50 µL high glucose DMEM media (ThermoFisher, Cat#11965092). The DNA mix was then added to Genjet mix to prepare transfection mix and incubated at room temperature for 15 minutes. 100 µL of the transfection mix was then added dropwise to each well containing 1 mL of complete media. Transfection media was removed and replaced with complete media after 5 hours. Surface markers were analyzed 28 hours post transfection as described previously (29).

### Transfection of BJAB cells

Transfections of BJAB cells with ctrl, WT-K5, or M-K5 expressing plasmids were performed using 4D-Nucleofector X Unit (Lonza), program SF at EN 150 as per the manufacturer’s protocol. Briefly, cells were washed with PBS and resuspended at 5×10^6^ cells per 100 µL SF nucleofection buffer. 1.25 µg of the DNA was then added to 100 µL nucleofection buffer and transferred to a cuvette for nucleofection. Nucleofected cells were incubated in the cuvettes for 10 minutes at room temperature, after which 500 µL prewarmed complete media was added and the cells then transferred to 6-well plates. Cells were cultured for 28 hours before performing surface marker analysis by flow cytometry as described previously (29)..

### RNA extraction and RT-qPCR

Total RNA was extracted with Direct-zol RNA miniprep kit with on-column DNase I digestion (Zymo Research #R2053). 0.5 to 1 μg of total RNA was used for reverse-transcription with ReverTra Ace qPCR RT master mix (Toyobo #FSQ-101) and quantitative PCR (qPCR) was performed with Thunderbird Next SYBR qPCR mix (Toyobo #QPX-201) and StepOnePlus real-time PCR system (ThermoFisher) following manufacturer’s instructions. Any targets with a Ct of <34 were labeled as “not detected” or “nd”. Copy level abundance was determined using standard curves generated using purified genomic stocks (KSHV bacterial artificial chromosome, BAC16). Absolute copy number of genomic stocks was determined using digital droplet PCR (Biorad QX600).

---

**Fig. S1:**
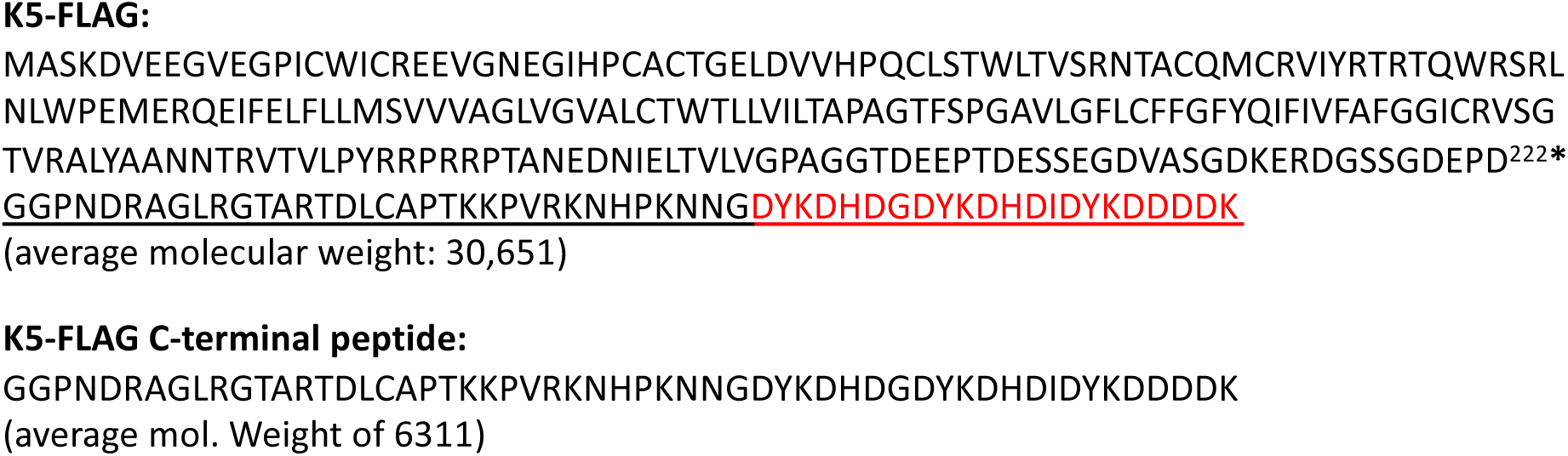
(A) Amino acid sequence for K5-FLAG expressed in BJAB cells. The 3XFlag sequence is shown in red. The predicted caspase cleavage site found using SitePrediction is indicated with an asterisk at D^222^*. (B) sequence of the predicted K5-FLAG peptide following caspase-cleavage at D222.

**Fig. S2:**
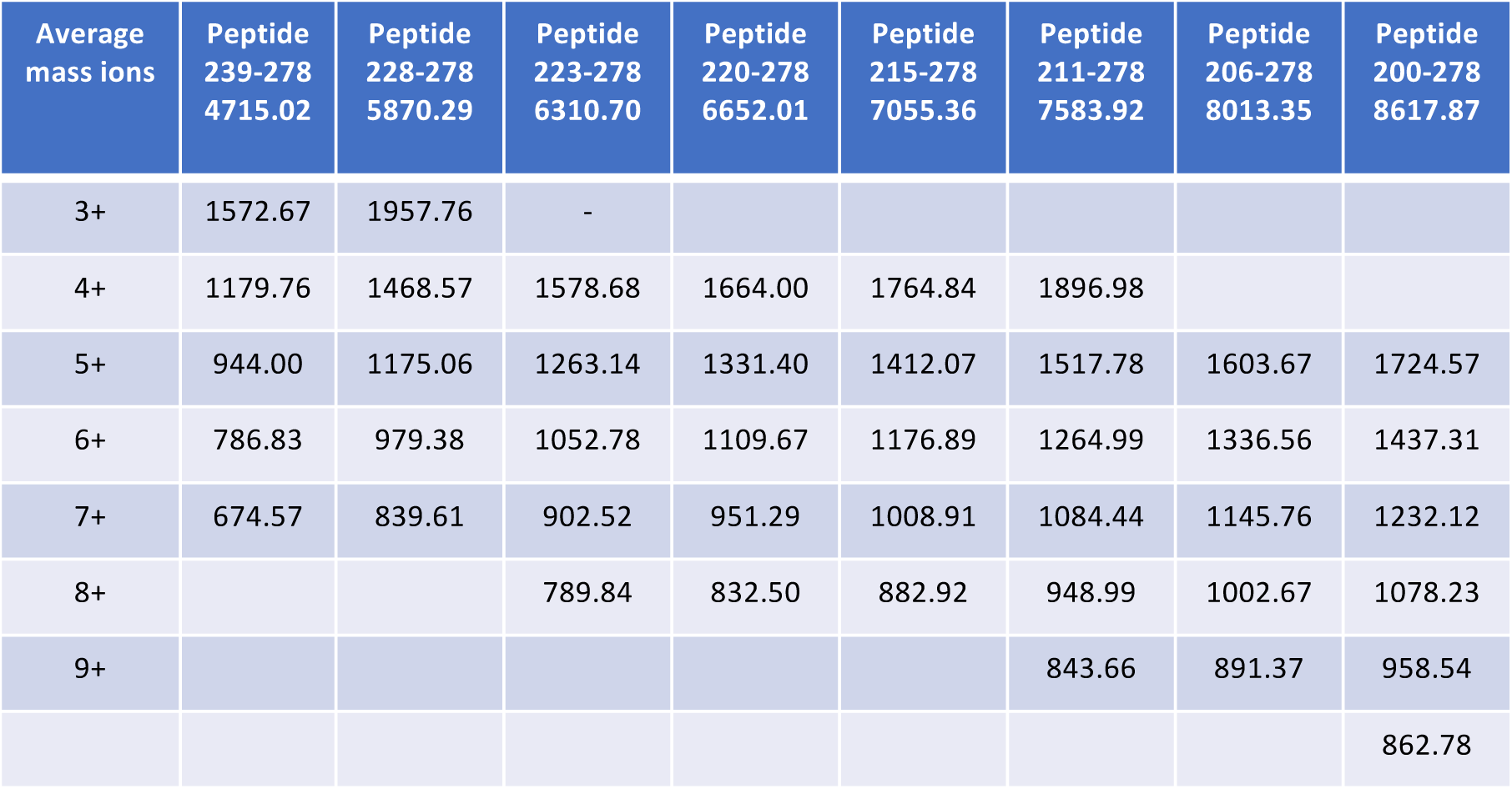
A. List of major ions expected for peptides generated by cleavage after aspartic acid. These ions were used to interrogate the eluted peptide from anti-FLAG beads. Only peptide ions for peptide cut at D222 (amino acids 223-278) were detected in the eluting peptides.

**Fig. S3:**
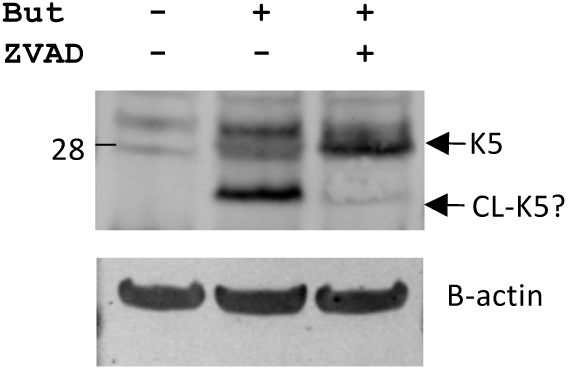
Caspase inhibition increases K5 protein levels in cytoplasmic extracts of JSC-1 cells. JSC-1 cytoplasmic protein extracts were analyzed by western blot using the Genescript rabbit K5 antibody. (Upper) Cytoplamsic protein lysates were extracted from JSC-1 cells after 48h of treatment with DMSO control and PBS control (lane 1), DMSO and 1 mM butyrate (lane 2), or ZVAD and butyrate (lane 3). (lower) B-actin shown as the loading control. CL-K5 is the presumed larger cleavage product.

**Fig. S4:**
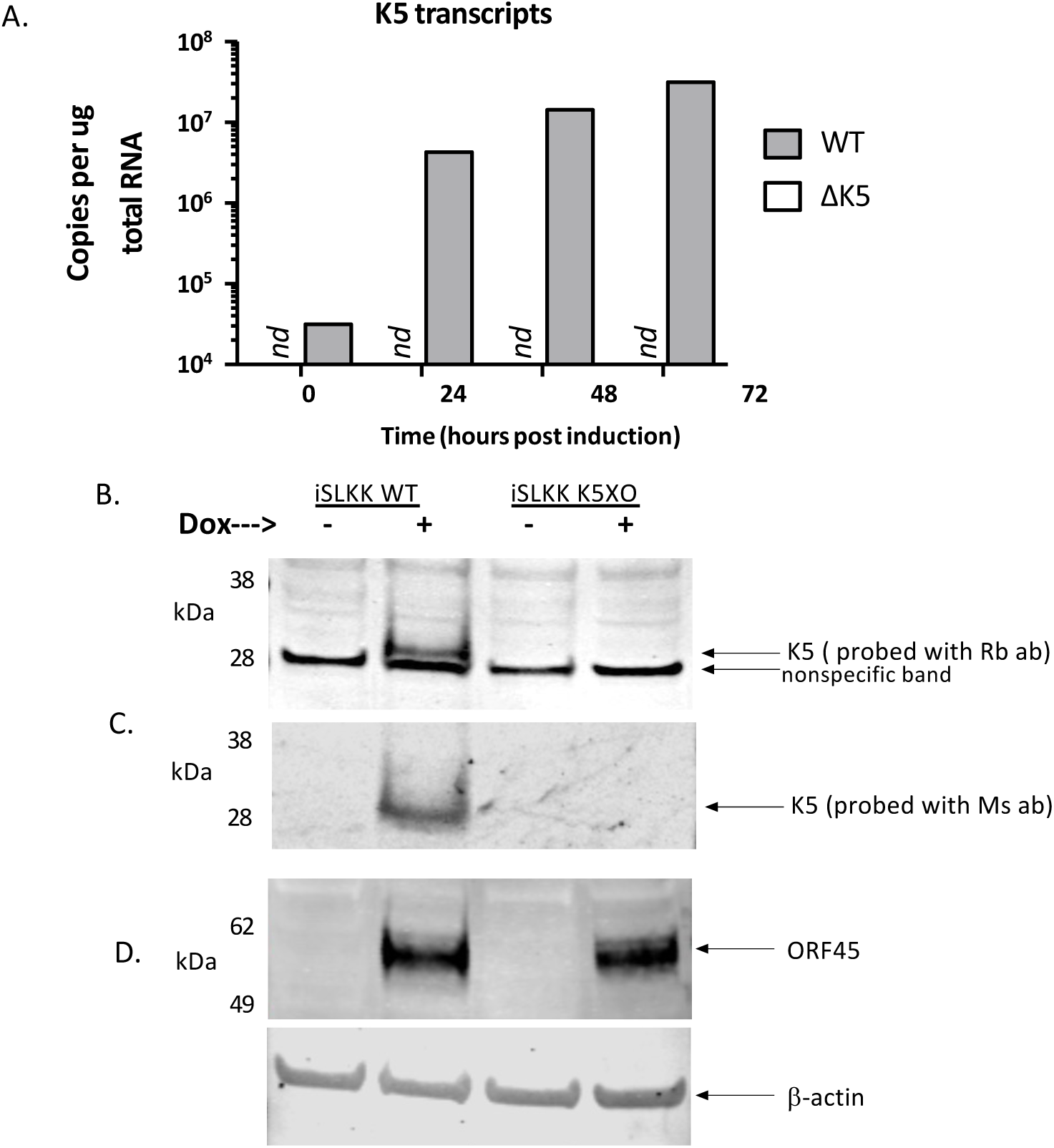
(A) qPCR on transcripts from WT iSLKK and delta-K5 iSLKK using a standard curve with known genome and KSHV BAC16 genome concentrations to quantify transcripts present (nd = ct > 34, nothing detected based on melt curve). K5 transcripts are detected in uninduced WT cells and increase upon Dox induction. No transcripts were detected in the delta-K5 iSLKK cells. Western blot of cytoplasmic lysates from uninduced and induced (Dox) WT and delta-K5 iSLKK cells for detection of K5 using the (B) rabbit (Rb) antibody or (C) mouse (Ms) antibody directed towards K5. (D) Western blot for detection of ORF45 lytic protein from the same lysates to confirm lytic induction in WT and K5 lines.

**Fig. S5:**
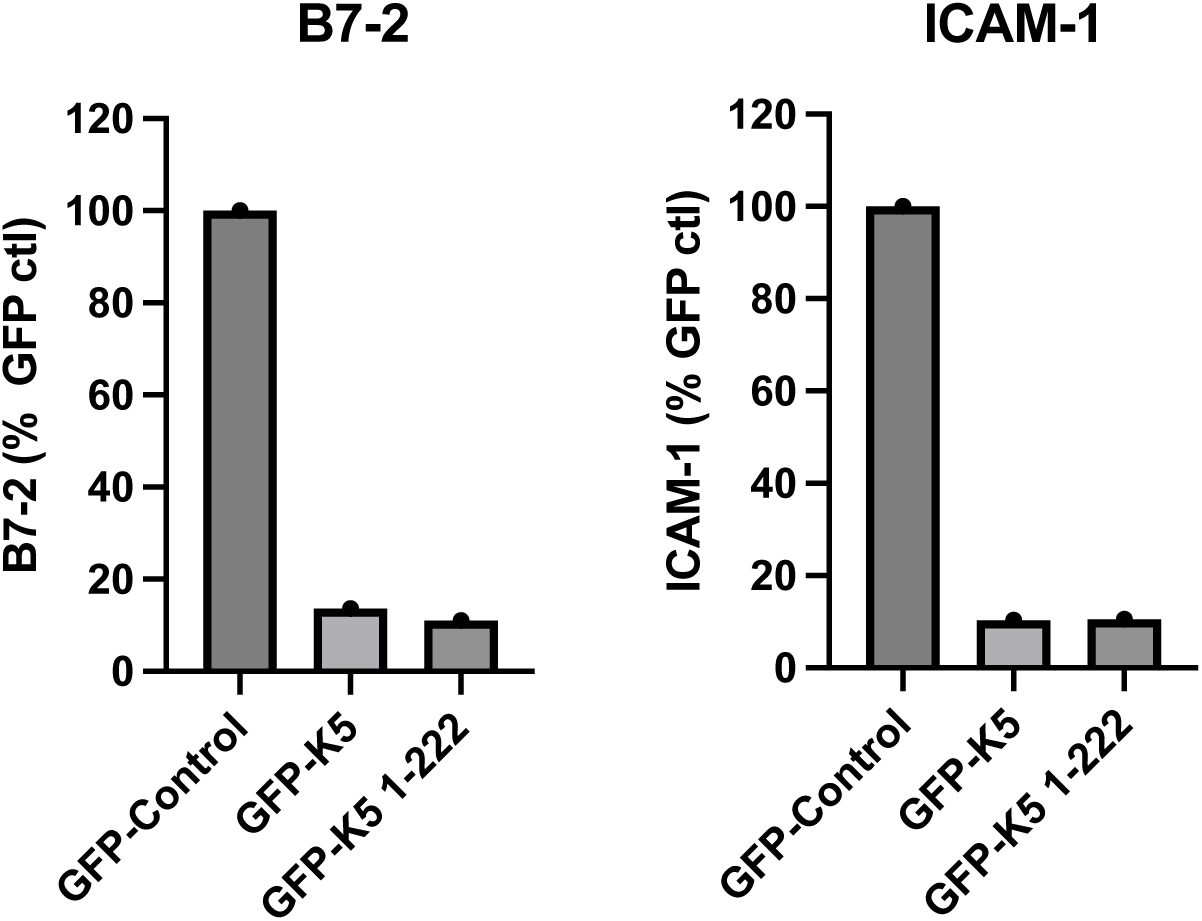
B7-2 and ICAM-1 surface expression in BJAB cells transfected with GFP-control vector, GFP-K5 plasmid, or truncated GFP-K5 (1–222) plasmid. Cells were transfected and then surface expression on live cells was measured (28h after transfection by FACS. Only one experiment was performed.

**Fig. S6:**
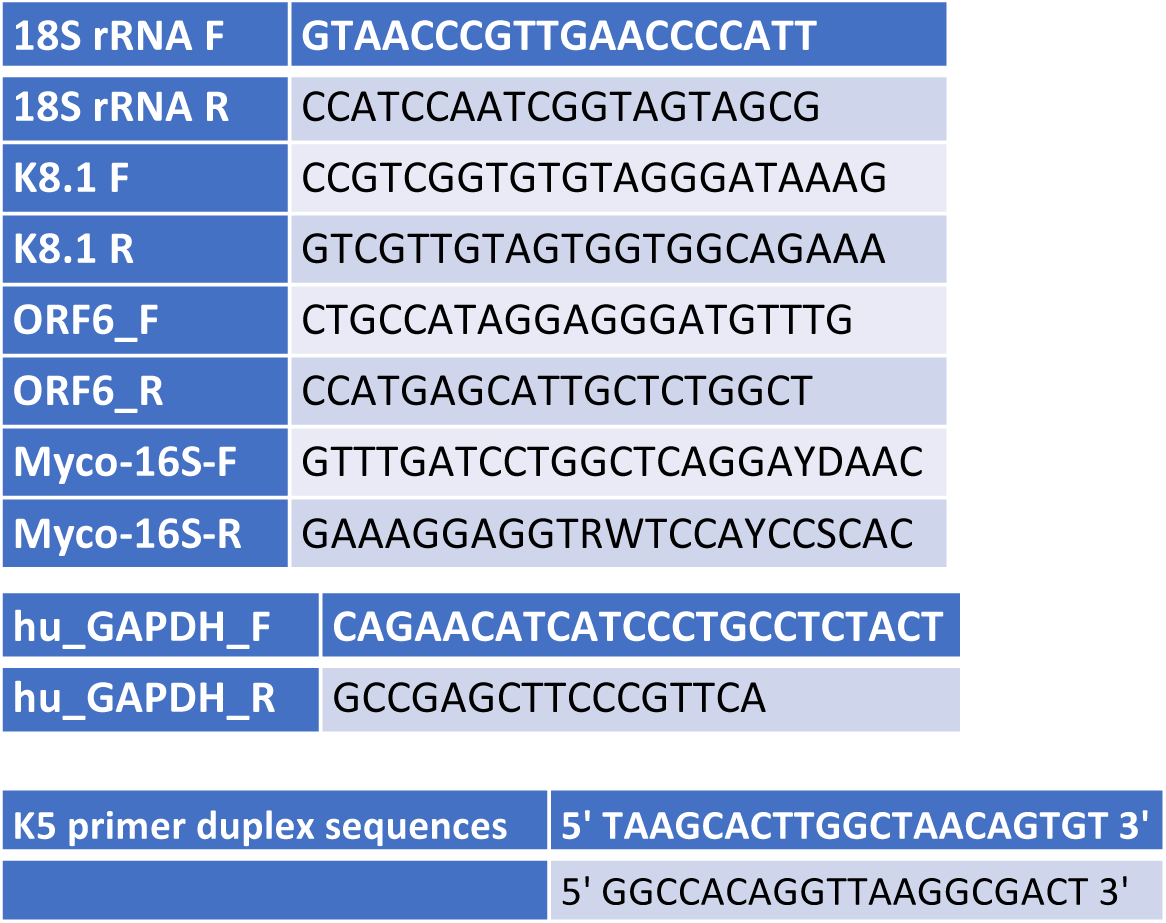
Forward and reverse primer sequences used in quantitative PCR analysis.

**Table 1:** KSHV protein sequences translated from the KSHV DNA sequence from BCBL-1 (accession number U93872.2) were analyzed for caspase cleavage sites for caspases-1,3,6,7, and 8 using site prediction and cross referenced with prosperousplus (http://prosperousplus.unimelb-biotools.cloud.edu.au/index.php/index). The proteins with sites scoring over 100 in SP are shown (35 of 87 proteins) along with the highest scoring caspase site for each protein. An * indicates that this protein has one or more other sites scoring over 100. The score along with the type of caspase (initiator caspase-8 (In), inflammatory caspase-1 (Inf), executioner caspases-3,6,7 (Ex) predicted to cleave at this site is shown. The prosperousplus score for the same SP sites are shown along with the PP predicted caspases to cleave the site. The final column indicates proteins known to be cleaved by caspases in KSHV-infected cells. Note the repeat region for LANA is excluded.

| KSHV Protein | Top scoring site by Site Prediction | SP score and predicted caspase class (I, Ex, Inf) | PP2 probability score and predicted caspase class (I, Ex, Inf) | Confirmed? |
| --- | --- | --- | --- | --- |
| ORF45* | DEED<sup>99</sup>*EDEE | 2787 (Ex) | 0.999 (I) | No |
| K10.5/vIRF3 | EEVD<sup>88</sup>*DGAG | 2679 (Ex) | 0.999 (Ex) | No |
| K5/MIR2* | DEPD<sup>222</sup>*GGPN | 1852 (I,Ex,Inf) | 1.000 (Ex, Inf) | This Study (I, Ex) |
| ORF57/MTA* | DETD<sup>33</sup>*APTL | 1742 (I,Ex, Inf) | 1.000 (I, Ex, Inf) | Yes (Ex) |
| ORF38 | VDVD<sup>18</sup>*GEPL | 1649 (Ex) | 0.988 (I, Inf) | No |
| ORF27* | VETD<sup>97</sup>*AI | 1146 (Ex) | 1.000 (I) | No |
| ORF22* | DESD<sup>648</sup>*GLQS | 984 (I, EX, Inf) | 1.000 (I, Inf) | No |
| ORF25* | VEGD<sup>346</sup>*KA | 926 (Ex) | 0.992 (Ex) | No |
| ORF10 | VEGD<sup>208</sup>*PE | 863 (Ex) | 0.977 (In) | No |
| ORF17* | VEKD<sup>518</sup>*APTP | 728 (Ex) | 0.999 (In) | No |
| ORF73 LANA* | DSVD<sup>53</sup>*GREC | 686 (I, Ex) | 1.000 (I, Ex, Inf) | Yes (Ex) |
| ORF63* | TEYD<sup>529</sup>*ED | 518 (Ex) | 0.970 (I) | No |
| ORF62 | VDLD<sup>208</sup>*ES | 504 (Ex) | 0.991 (Ex) | No |
| ORF70 | VDAD<sup>174</sup>*AD | 487 (Ex) | 0.996 (I) | No |
| ORF59 | ESPD<sup>332</sup>*SPPL | 464 (Inf) | 0.990 (Inf) | No |
| ORF67 | VESD<sup>116</sup>*VY | 424 (Ex) | 0.998 (Ex) | No |
| ORF37* | DTLD<sup>21</sup>*GLTV | 413 (Ex) | 0.999 (Ex) | No |
| ORF64* | DESD<sup>1966</sup>*TASG | 392 (Ex) | 1.000 (Ex) | No |
| K8* | IEED<sup>28</sup>*LS | 320 (I, Ex) | 0.998 (I) | No |
| ORF54 | GETD<sup>93</sup>*KD | 297 (Ex) | 0.985 (8) | No |
| ORF50/RTA | DSPD<sup>408</sup>*NPSS | 276 (Inf) | 0.998 (Ex) | No |
| K11/vIRF2* | LAPD<sup>27</sup>*SPRP | 274 (Inf) | 0.981 (I) | No |
| ORF6* | TEED<sup>1039</sup>*VI | 262 (Ex) | 0.998 (Inf) | No |
| ORF34 | EAVD<sup>139</sup>*GLCD | 256 (Ex) | 1.000 (Inf) | No |
| ORF21/TK* | TDDD<sup>116</sup>*SG | 253 (Ex) | 0.988 (I) | No |
| ORF75* | IEDD<sup>184</sup>*VI | 252 (Ex) | 1.000 (Inf) | No |
| ORF71/vFLIP | TDVD*ALMS | 214 (Ex) | 0.999 (I) | No |
| K3/MIR1* | EDED<sup>5</sup>*VP | 183 (Ex) | 0.929 (Ex) | No |
| ORF4 | SLTD<sup>495</sup>*SA | 180 (I, Ex, Inf) | 1.000 (Ex) | No |
| ORF19 | LELD<sup>71</sup>*RL | 153 (Ex) | 0.989 (Ex) | No |
| ORF20 | EVLD<sup>30</sup>*SSSE | 135 (Ex) | 1.000 (Inf) | No |
| K14* | TDSD<sup>178</sup>*GLTV | 127 (Ex) | 0.997 (I) | No |
| ORF74/GPCR | LDDD<sup>14</sup>*ES | 125 (Ex) | 0.899 (I) | No |
| ORF2 | SSLD<sup>94</sup>*AALG | 115 (Ex) | 0.996 (Ex) | No |
| ORF49 | QELD<sup>31</sup>*TL | 101 (Ex) | 0.998 (Ex) | No |

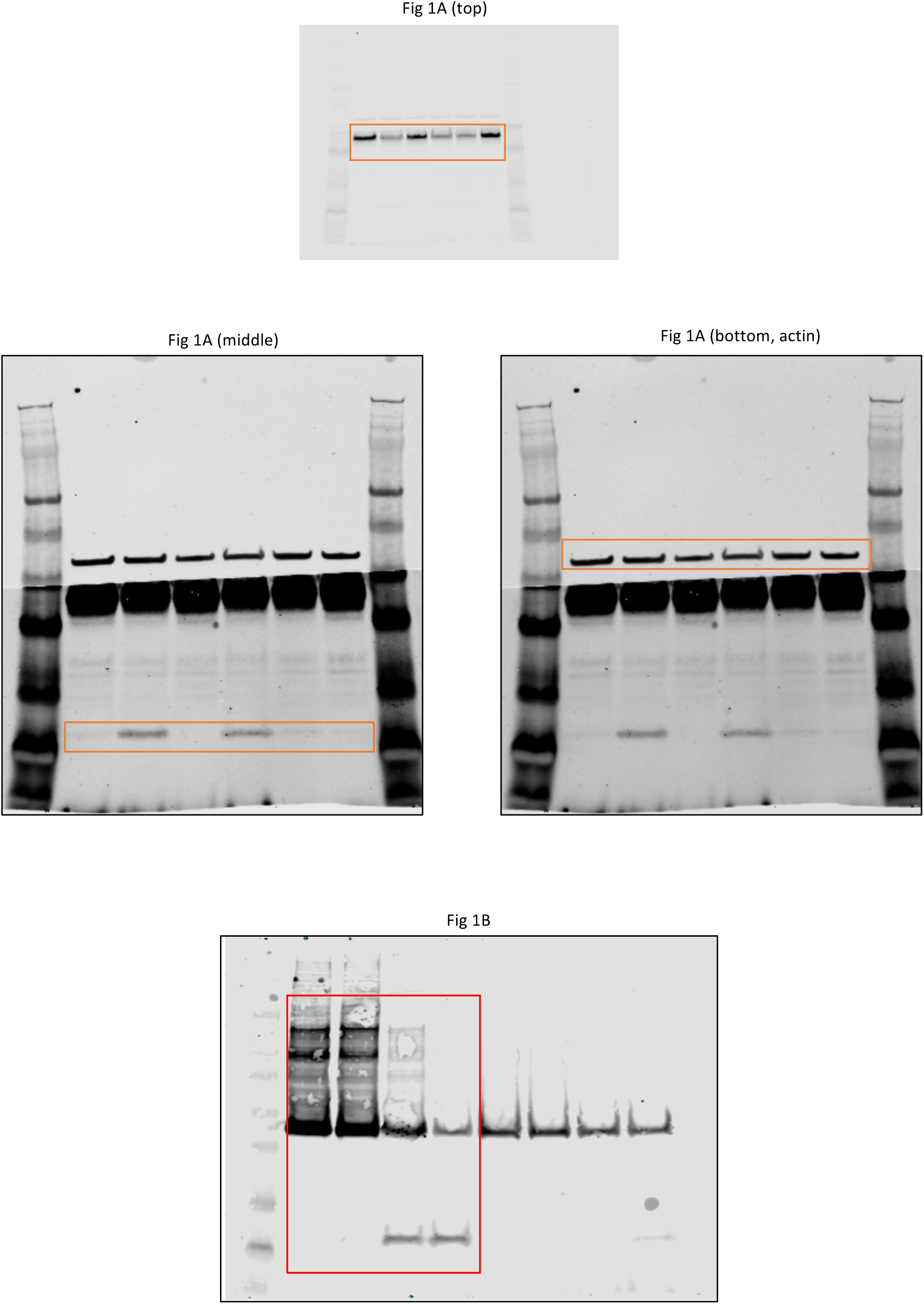
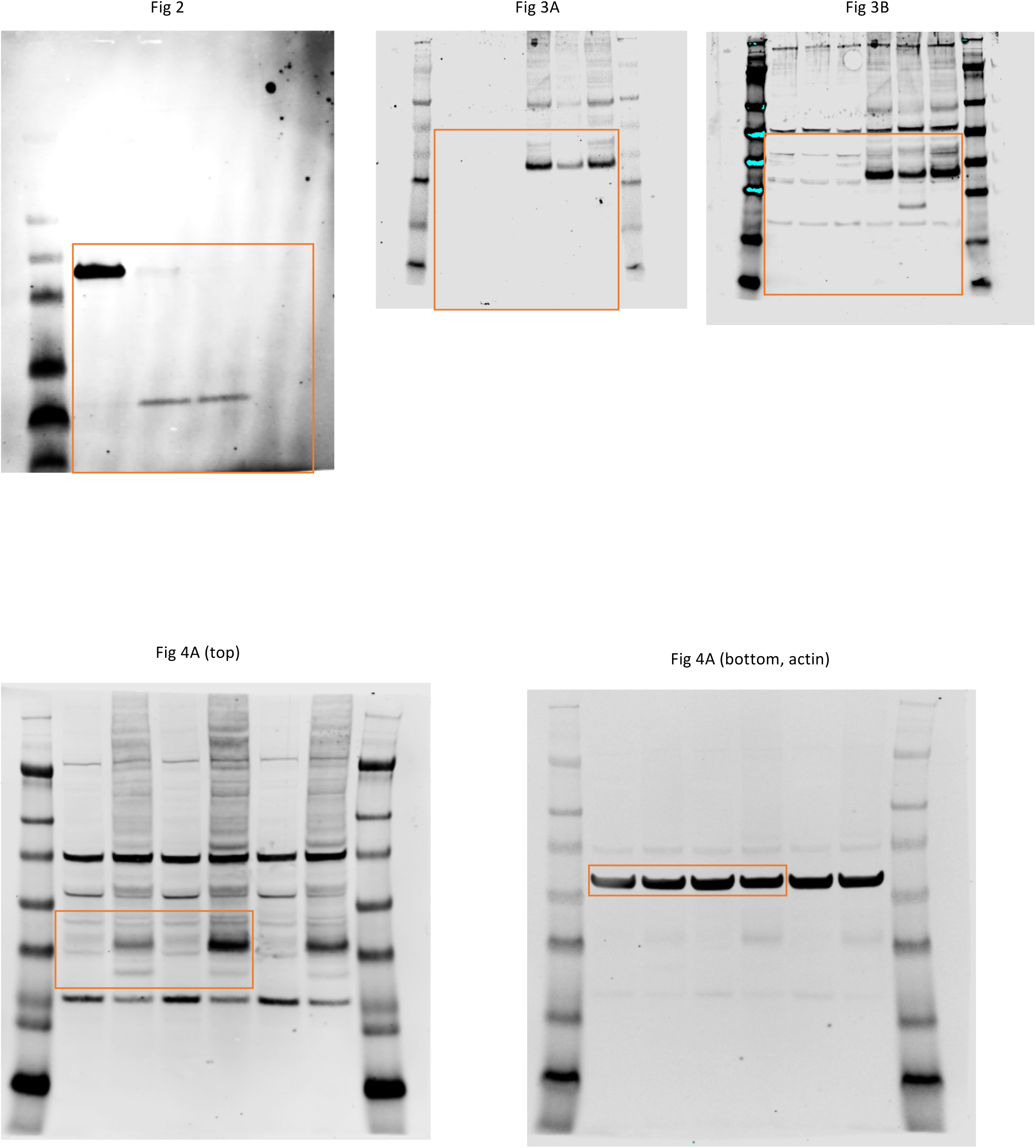
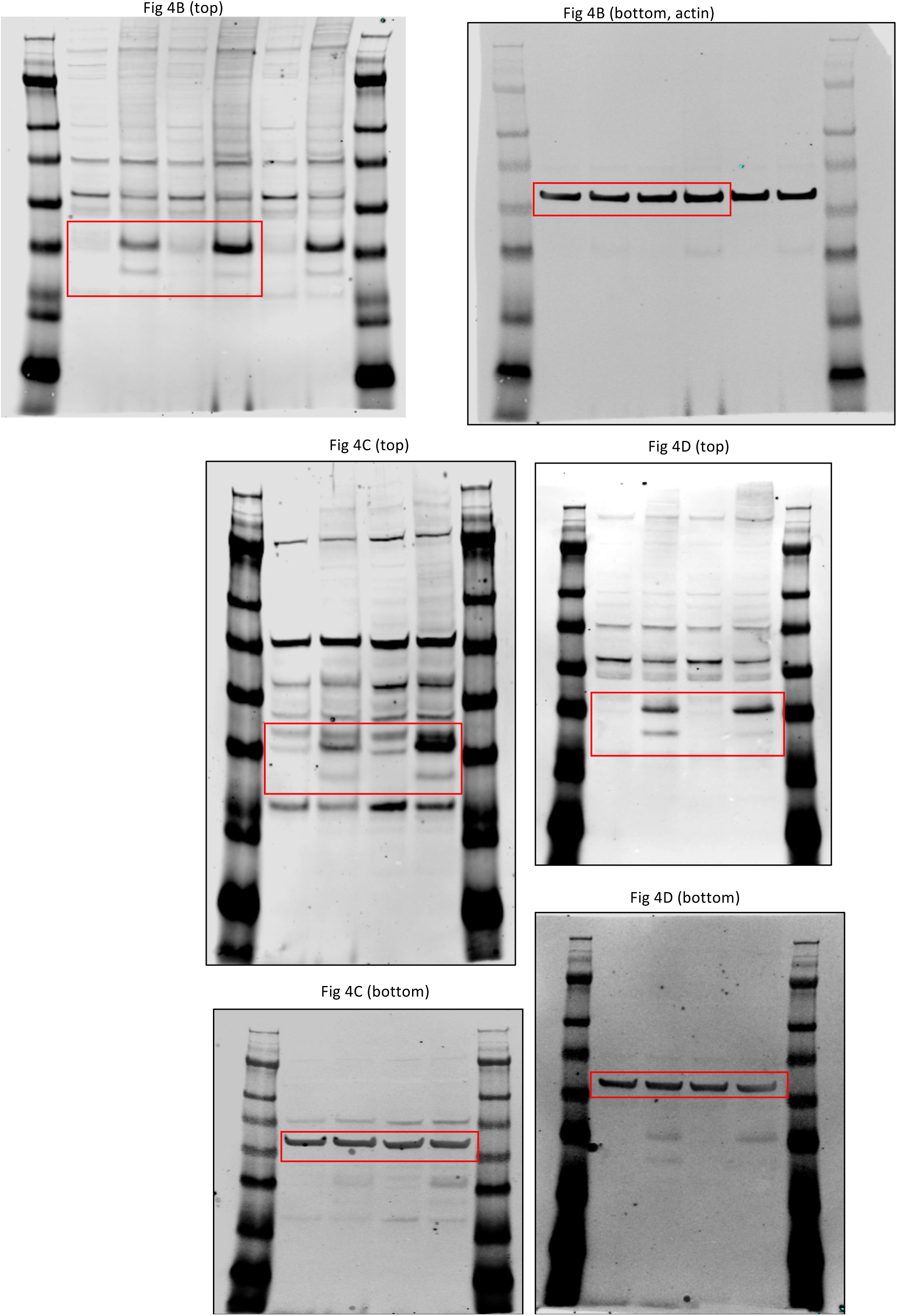
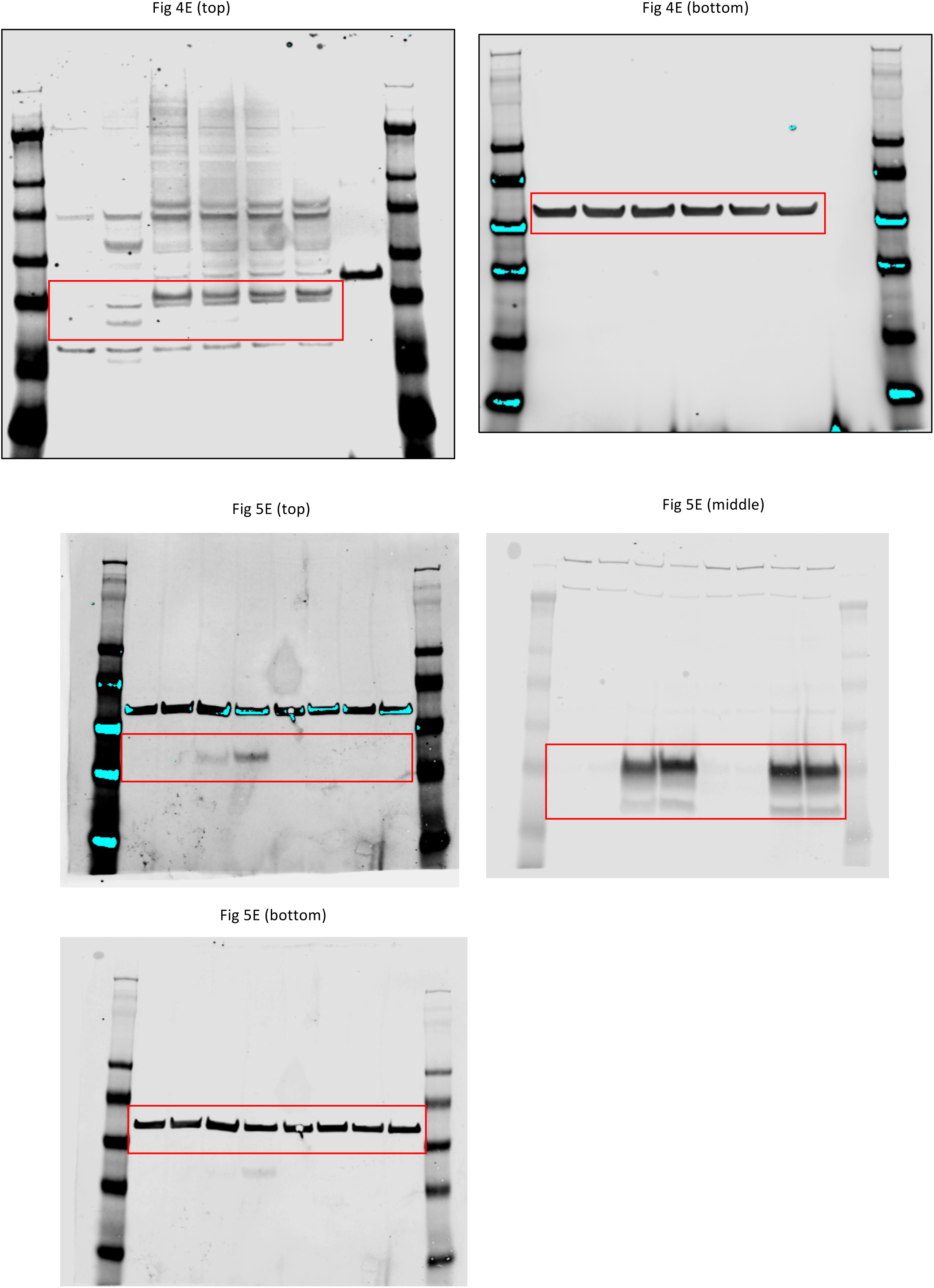
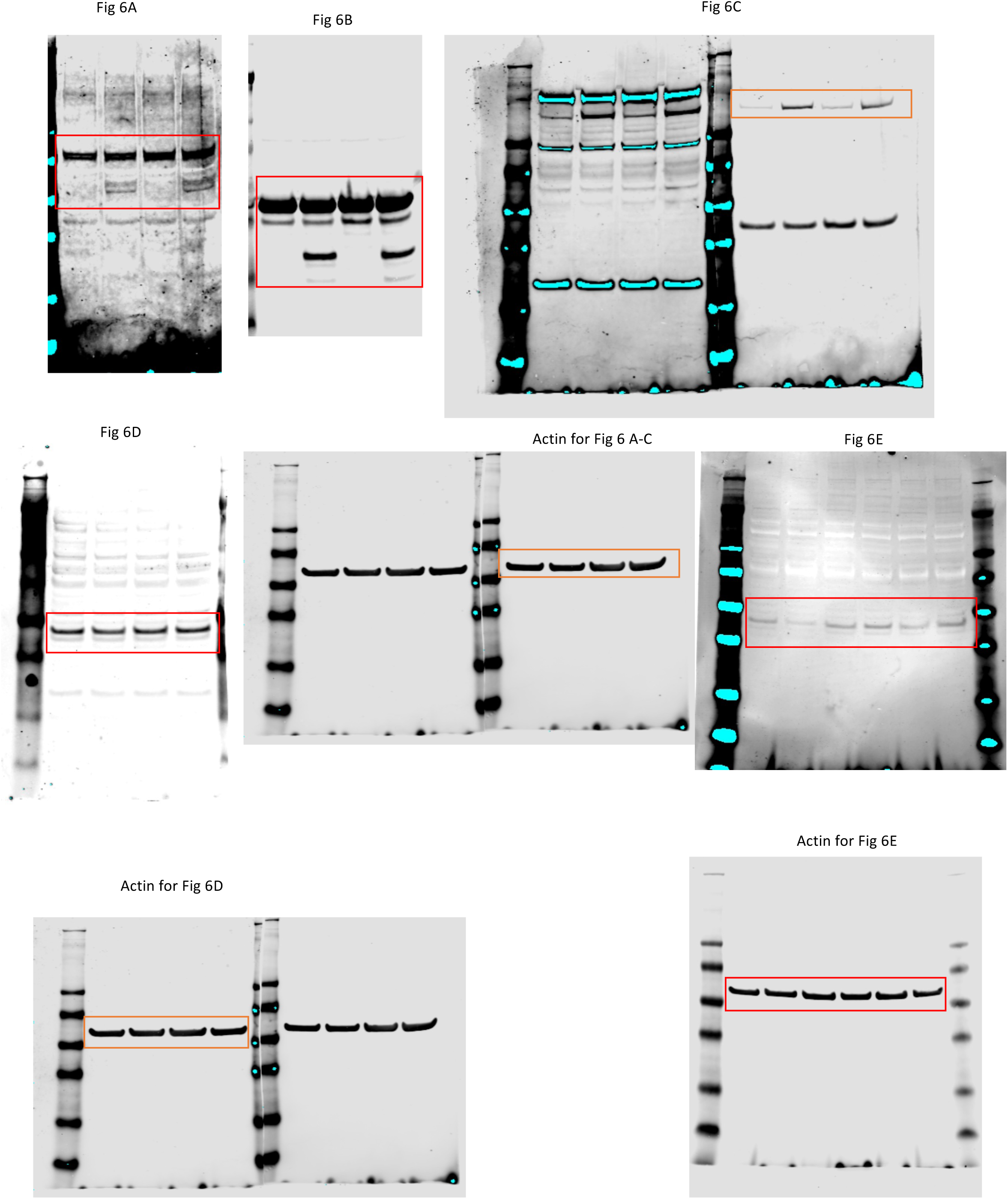
Full scans for immunoblots.

